# A plastidial DEAD box RNA helicase plays a critical role in high light acclimation by modulating ribosome biogenesis in *Chlamydomonas reinhardtii*

**DOI:** 10.1101/2022.05.16.492170

**Authors:** El Batoul Djouani-Tahri, Sreedhar Nellaepalli, Pascaline Auroy, Emmanuelle Billon, Adrien Burlacot, Frédéric Chaux-Jukic, Stéphan Cuiné, Virginie Epting, Marie Huleux, Bart Ghysels, Miriam Schulz-Raffelt, Isabelle Te, Sabine Brugière, Yohann Couté, Yuichiro Takahashi, Yonghua Li-Beisson, Gilles Peltier

## Abstract

Photosynthetic organisms have developed sophisticated strategies to fine-tune light energy conversion to meet the metabolic demand, thereby optimizing growth in fluctuating light environments. Although mechanisms such as energy dissipation, photosynthetic control, or the photosystem II (PSII) damage and repair have been widely studied, little is known about the regulation of protein synthesis capacity during light acclimation. By screening a *Chlamydomonas reinhardtii* insertional mutant library using chlorophyll fluorescence imaging, we isolated a high chlorophyll fluorescence mutant (*hf_0_*) defected in a gene encoding a putative plastid targeted DEAD-box RNA helicase called CreRH22. CreRH22 is rapidly induced upon illumination and belongs to the GreenCut, a set of proteins specific to photosynthetic organisms. While photosynthesis is slightly affected in the mutant under low light (LL), exposure to high light (HL) induces a marked decrease in both PSII and PSI, and a strong alteration of the light-induced gene expression pattern. These effects are explained by the inability of *hf_0_* to increase plastid ribosome amounts under HL. We conclude that CreRH22, by promoting ribosomal RNA precursor maturation in a light-dependent manner, enables the assembly of extra-ribosomes required to synthesize photosystem subunits at a higher rate, a critical step in the acclimation of algae to HL.

## INTRODUCTION

In natural environments, photosynthetic organisms face highly fluctuating light conditions, and developed a complex network of regulatory mechanisms to adjust the capture and transformation of light energy to the metabolic demand. Such mechanisms are highly desirable since any disequilibrium between the conversion and use of light energy may result in severe photo-damage due to the formation of harmful reactive oxygen species (Niyogi, 1999). A wide range of mechanisms dynamically regulate light-energy conversion and photosynthetic electron flow, and operate at different time scales (Erickson et al., 2015). The short-term response involves energy dissipation by photosystems through a pH-dependent de-excitation of chlorophylls (Peers et al., 2009). On longer time scales (minutes to hour), other acclimation mechanisms operate in addition to the pH dependent NPQ, including state transitions (Depege et al., 2003; Allorent et al., 2013), adjustment of chlorophyll antenna size and photosystem stoichiometry (Bonente et al., 2012; Erickson et al., 2015). The PSII damage and repair cycle, although initially considered as a wasteful process (called photoinhibition) and involving selective destruction of PSII core subunits followed by active protein synthesis (Ohad et al., 1984), is now viewed as part of a dynamic and sophisticated machinery in which PSII is degraded and further repaired (Murata et al., 2012; Adams et al., 2013; Erickson et al., 2015; Li et al., 2018). The PSII repair cycle would protect the photosynthetic electron transfer chain, particularly PSI (Tikkanen et al., 2014). During the PSII repair cycle, translation of plastid-encoded PSII subunits D1, D2 and CP43 is enhanced (Christopher and Mullet, 1994; Kettunen et al., 1997; Minai et al., 2006; Jarvi et al., 2015). However, despite extensive studies on PSII repair, mechanisms involved in the regulation of translation remain largely unknown (Sun and Zerges, 2015). Analysis of gene expression during a day-night cycle showed that nuclear genes encoding chloroplast ribosome proteins are rapidly and transiently expressed upon sudden HL exposure, indicating that ribosome biogenesis is critical in such conditions (Zones et al., 2015).

DEAD-box RNA helicases catalyze ATP-dependent unwinding of RNA structures, and are involved in many aspects of RNA metabolism, including RNA synthesis, cleavage, modification, or ribosome biogenesis (Silverman et al., 2003). *CrhR*, the unique DEAD-box RNA helicase present in the genome of the cyanobacterium *Synechocystis* sp. PCC 6803, participates in redox regulation of gene expression (Kujat and Owttrim, 2000), low temperature acclimation of photosynthesis (Rosana et al., 2012; Sireesha et al., 2012), and was recently shown to interact with transcripts associated with photosynthesis (Migur et al., 2021). In plant chloroplasts, DEAD-box RNA helicases participate in ribosome biogenesis (Asakura et al., 2012; Chi et al., 2012), chloroplast differentiation (Wang et al., 2000), and responses to diverse abiotic stresses (Nawaz and Kang, 2017), but a role of these enzymes in the light-response of eukaryotic photosynthesis has not been reported so far.

With the aim to unveil new regulatory and acclimation mechanisms of photosynthesis, a forward genetic approach was developed in the unicellular microalga *Chlamydomonas reinhardtii* by screening an insertional mutant library based on the analysis of chlorophyll fluorescence (Tolleter et al., 2011). We report here on the isolation of a high chlorophyll fluorescence mutant (*hf_0_*) defected in a gene encoding a putative plastid DEAD-box RNA helicase (*CreRH22*), and show its involvement in ribosomal RNA processing and ribosome biogenesis. *CreRH22* belongs to the GreeenCut2 (Karpowicz et al., 2011) and its expression is rapidly and transiently induced upon HL exposure. Photosynthetic activity and photoautotrophic growth of the *hf_0_* mutant are affected at HL intensity. We conclude that rapid induction of CreRH22 promotes ribosome biogenesis upon HL exposure, thus increasing the translation capacity and enabling efficient turnover of plastid-encoded photosystems subunits.

## RESULTS

### Isolation of high chlorophyll fluorescence *Chlamydomonas* mutant defected in a DEAD-box RNA helicase

From the screening of an insertion mutant library based on the analysis of chlorophyll fluorescence transients of *Chlamydomonas* colonies grown in photo-autotrophic conditions (Tolleter et al., 2011), a mutant (*hf_0_*) showing a high F_0_ fluorescence level was isolated (Fig. 1A). The *hf_0_* mutant harbors a single insertion of the *AphVIII* paromomycin resistance cassette (Fig. 1B and Supplemental Fig. S1) located within the 7^th^ exon of a gene (Cre03.g166650) encoding a putative DEAD-box RNA helicase named as CreRH22 (Fig. 1B). The Cre03.g166650 transcript was not detected by RT-PCR in *hf_0_* (Fig.1C). CreRH22 shares common properties of DEAD-box RNA helicases, including the Asp-Glu-Ala-Asp (DEAD) sequence, an RNA-binding motif and an ATP hydrolyzing domain (Fig. 1B). Phylogenetic analysis showed that *CreRH2* belongs to a DEAD-box RNA helicase clade containing a previously characterized member in Arabidopsis (AtRH22) (Chi et al., 2012) (Supplemental Fig. S2). AtRH22 was reported to be targeted to the chloroplast and to be involved in ribosomal RNA processing and chloroplast ribosome biogenesis (Chi et al., 2012). By using the prediction tool Predalgo specifically developed for intracellular targeting of microalgae (Tardif et al., 2012), CreRH22 was predicted to be chloroplast targeted, like six other putative DEAD-box RNA helicases encoded by the *C. reinhardtii* genome (Supplemental Fig. S3). Moreover, the CreRH22 protein belongs to the GreenCut2 (identified as Hel15), an inventory of proteins identified on a phylogenomic basis, conserved in plants and green algae but not in non-photosynthetic organisms (Karpowicz et al., 2011). Complementation was attempted by both nuclear and chloroplast transformation. For nuclear complementation, a genomic DNA fragment corresponding to the full *CreRH22* gene was cloned into a vector containing a hygromycin resistance cassette, and resistant clones were screened by chlorophyll fluorescence imaging (Supplemental Fig. S4). Full complementation was observed in one strain out of 400 hygromycin resistant clones. For plastid transformation, a synthetic gene with a codon usage adapted to the *Chlamydomonas* chloroplast genome was used (Supplemental Fig. S5). Two independent homoplasmic chloroplast transformants were analyzed, chlorophyll fluorescence measurements showing a full recovery of the wild-type phenotype in both transformants (Supplemental Fig. S4). We conclude that inactivation of *CreRH22* is responsible for the *hf_0_*chlorophyll fluorescence phenotype, complementation by plastid expression confirming chloroplast targeting.

**Figure 1.**
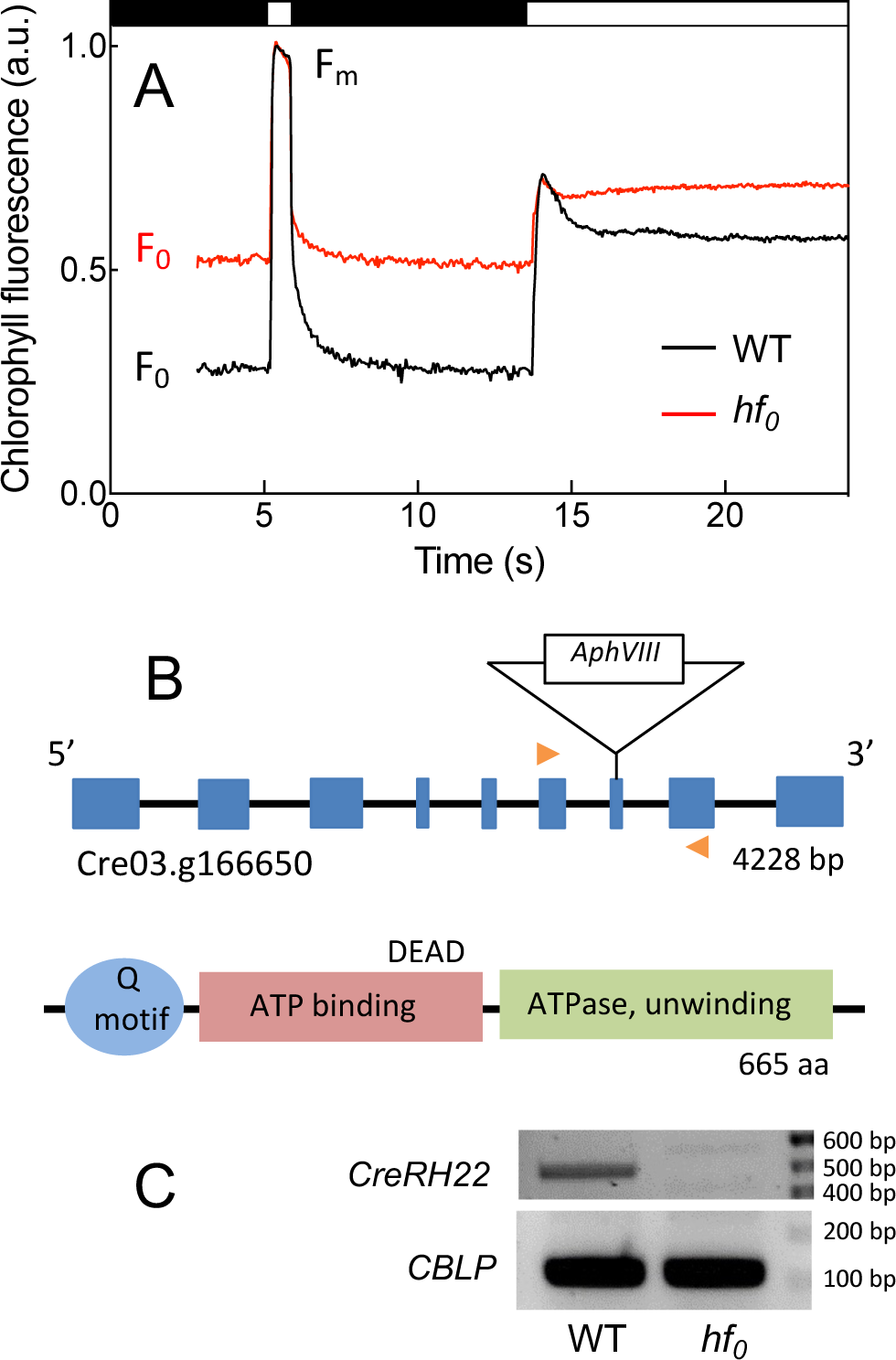
Isolation and molecular characterization of *hf_0_*, a *C. reinhardtii* high chlorophyll fluorescence mutant. **(A)** The *hf_0_* mutant was isolated from the screening of an insertion library based on the analysis of chlorophyll fluorescence transients (after recording F_0_ fluorescence in the dark, a saturating flash at 5s was followed by 240 µmol photons m^-2^ s^-1^ actinic red light at 14 s). **(B)** The paromomycin resistance cassette (*AphVIII* gene) is integrated in exon 7 of the Cre03.g166650 locus encoding a gene annotated as a DEAD-box RNA helicase, renamed here *CreRH22* based on a phylogenetic analysis (Supplemental Fig. 2). The *CreRH22* gene is composed of 9 exons (blue boxes) and 8 introns (black lines). Orange arrows show primers used for RT-qPCR shown in (C). The CreRH22 protein shows typical features of DEAD-box RNA helicases, including a Q motif, an ATP binding domain containing a DEAD motif, and an ATPase RNA unwinding helicase domain. **(C)** RT-qPCR showing the absence of the *CreRH22* gene transcript in *hf_0_*. *CBLP* (alias RACK1 Cre06.g27822) is used as loading control.

We observed that the chlorophyll fluorescence phenotype of the *hf_0_*mutant vanishes when batch cultures reached a high cell density. Since the photon flux received by each cell is lower in a dense culture than in a diluted culture due to cell-shading, we hypothesized that the mutant phenotype may depend on the light intensity. Indeed, when cells were grown under HL (240 µmol photons. m^-2^ s^-1^) chlorophyll fluorescence induction of the mutant was strongly affected, showing higher F_0_ and Fs levels and reduced PSII yields as compared to wild-type control and two complemented lines (Fig. 2A, C). However, when cells were grown under LL (15 µmol photons m^-2^ s^-1^) differences in the F_0_ levels and PSII yields between *hf_0_* and control strains were much smaller (Fig. 2B, D).

**Figure 2.**
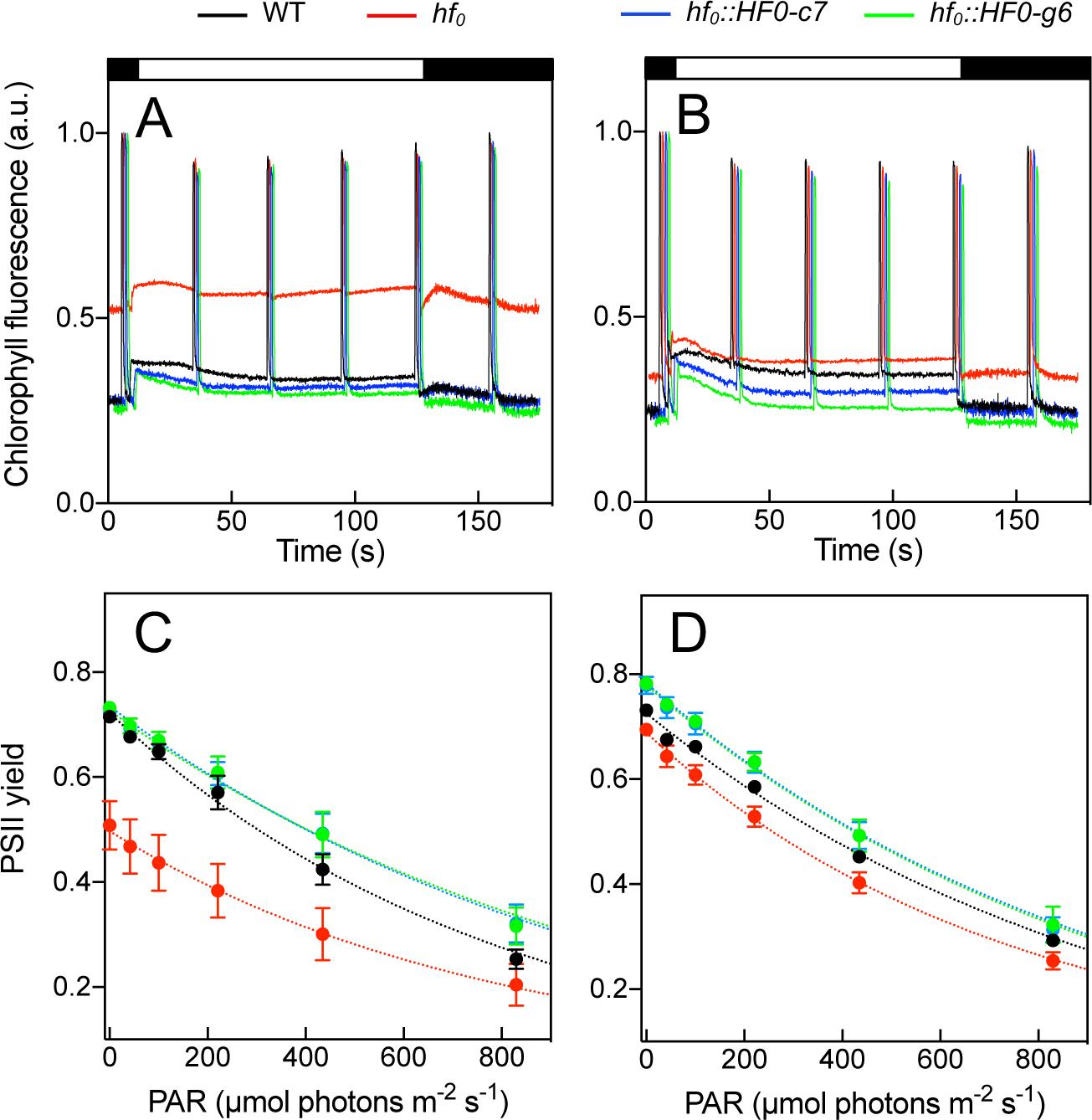
Effect of light on photosynthetic properties of the *hf_0_* mutant. Wild-type (WT), *hf_0_* and two complemented lines (*hf_0_::HF_0_-c7* and *hf_0_::HF_0_-g6*) were grown photo-autotrophically in 3% CO_2_-enriched air under HL (240 µmol photons m^-2^ s^-1^ **A, C**) or LL (15 µmol photons m^-2^ s^-1^ **B, D**). Representative chlorophyll fluorescence measurements during a dark/light (100 µmol photons m^-2^ s^-1^) transient for HL **(A)** or LL **(B)** grown cells. Black and white boxes indicate dark and light periods, respectively. **(C, D)** PSII yields determined from chlorophyll fluorescence measurements as F_v_/F_m_ in the dark after 15 min dark adaptation or (F_m’_-F_s_)/F_m’_ in the light in HL **(C)** or LL **(D)** grown cells. Shown are means +/-SD (n=3).

### Ribosomal RNA processing is affected in *hf_0_*

We next investigated whether CreRH22 is involved in chloroplast ribosomal RNA processing by analyzing this mechanism in *hf_0_*. Chloroplast ribosomal RNAs are organized in polycistronic transcription units, which need to be matured into a set of overlapping RNAs through a series of processing steps. The non-mature polycistronic rRNA undergoes endo-nucleolytic cleavages to produce mature 7S, 3S, 23S and 5S rRNAs (Holloway and Herrin, 1998) (Fig. 3A). Accumulation and processing of chloroplast ribosomal RNAs were further examined by Northern blot using specific probes for mature 3S, 7S, and 23S rRNAs. We found that RNA precursor levels are increased in *hf_0_* as compared to the wild-type, mature RNA levels being reduced accordingly (Fig. 3B). Plastid transcript level encoding PSII and PSI subunits, analyzed by RT-qPCR, showed a slightly higher level of transcripts in *hf_0_* compared to the wild-type both under LL (30 µmol photons m^-2^ s^-1^) and HL (240 µmol photons m^-2^ s^-1^) (Supplemental Fig. S6A, B). Analysis of chloroplast transcripts encoding non-polycistronic ribosomal components (*rpl36* and *rps18)* showed no obvious difference in the levels of transcripts (Supplemental Fig. S6C, D), the 16S transcript level being lower in *hf_0_* compared to the wild-type (Supplemental Fig. S6E). We conclude from these experiments that, as previously reported for AtRH22 (Chi et al., 2012), CreRH22 is involved in chloroplast ribosomal RNA processing, its defect resulting in an accumulation of unprocessed rRNA forms and thereby a decrease in the amount of plastid ribosomes.

**Figure 3.**
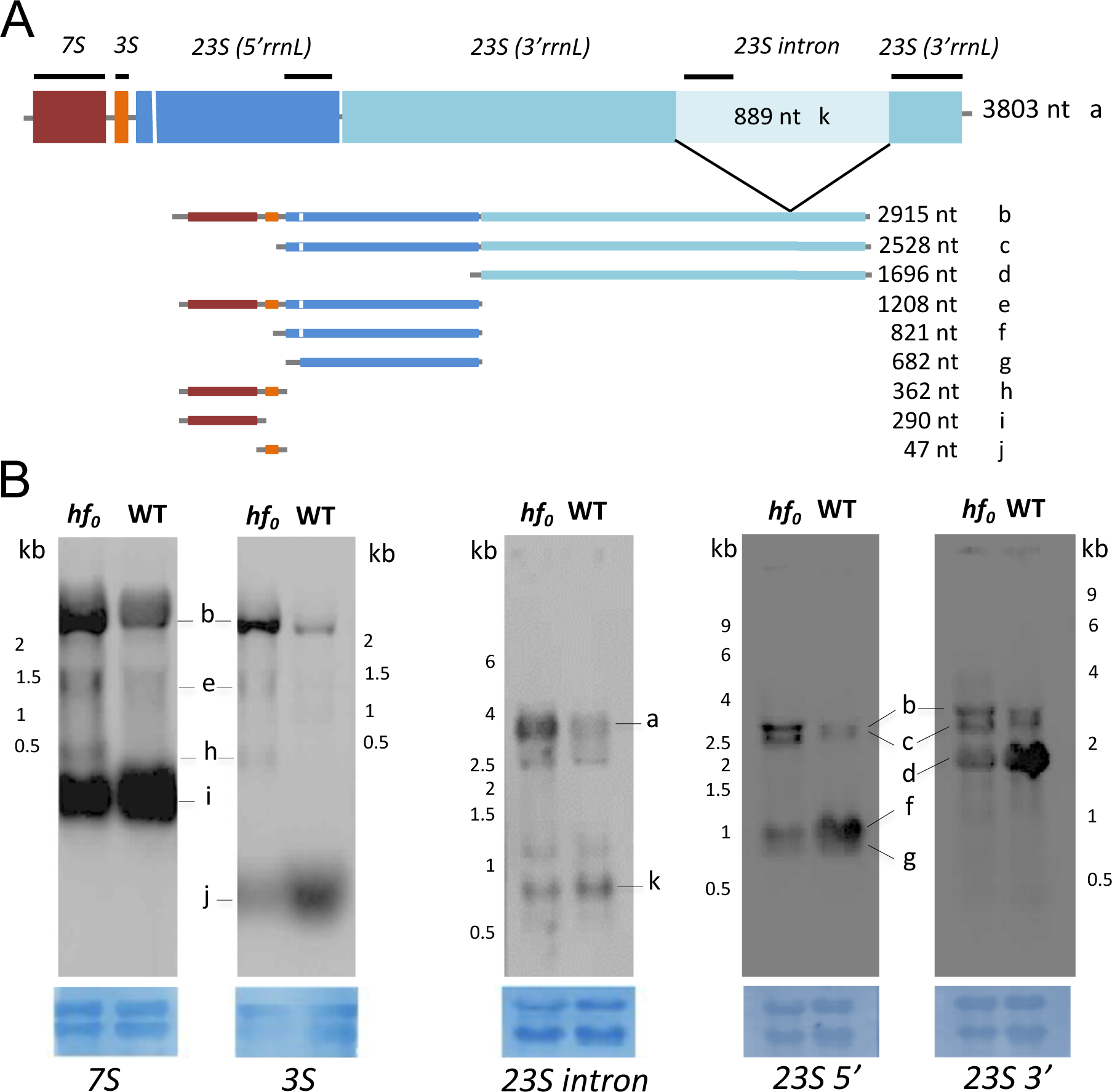
Ribosomal RNAs processing is altered in the *hf_0_* mutant. **(A)** Scheme showing the organization of the *C. reinhardtii* chloroplast ribosomal RNA (rRNA) operon and location of the different probes (black bars) used for Northern blot analysis. **(B)** rRNAs levels analyzed by Northern blot in WT and *hf_0_* cells grown photo-autotrophically under a light intensity of 60 µmol photons m^-2^ s^-1^. 5μg of total RNA were loaded on agarose gels, blotted on nylon membrane and hybridized with DIG-labeled probes. For loading controls (lower blue boxes), nylon membranes were stained with methylene blue after hybridization.

### Accumulation and turnover of PSII and PSI subunits are affected in *hf_0_*

To better characterize the effect of CreRH22 on protein synthesis, quantitative immuno-analysis was carried out on a set of chloroplast proteins, including plastid ribosomal proteins and components of the photosynthetic electron transport chain (Fig. 4). Under LL, no differences in protein amounts were observed between *hf_0_*and the wild-type, except for chloroplast ribosomal proteins L30 and S21, both present at lower amounts in *hf_0_*. Upon HL exposure, the amount of both ribosomal proteins increased in the wild-type while remaining at a low level in *hf_0_*. In *hf_0_*, HL exposure induced a marked decrease in the amounts of PSII subunits PsbA (D1), PsbC (CP43), and PSI subunits PsaC, PSAD and PSAF, this decrease being not observed in wild-type cells. The amounts of other chloroplast components, including AtpB, a subunit of the ATP synthase (CF0-CF1), and PetA a subunit of the cytochrome *b_6_f* complex. ^35^S pulse-labeling experiments of thylakoid proteins were then carried out to determine to what extent such changes in protein amounts reflect changes in protein synthesis activity. No difference in pulse labeling patterns was observed under LL between wild-type, *hf_0_* and two complemented lines (Fig. 5). However, upon 6h of HL exposure a lower D1 labeling was observed in *hf_0_*, the effect being reversed in both complemented lines. The turnover of D1 is known as the highest of plastid proteins due to the existence of an efficient damage/repair system (Jarvi et al., 2015). Note that slight differences in labeling between wild-type and *hf_0_*were also observed, although at a lower extent, on other proteins including LHCII and the PSI subunit PsaC. On the other hand, the labeling of ATP synthase (CF0-CF1) subunits AtpA and AtpB was unaffected in the mutant. We conclude from these experiments that the protein translation/synthesizing capacity in *hf_0_*is lower than in the wild-type due to a lower ribosome content, which becomes limiting under HL, and results in a decreased accumulation of plastid proteins actively synthesized under HL, such as D1.

**Figure 4.**
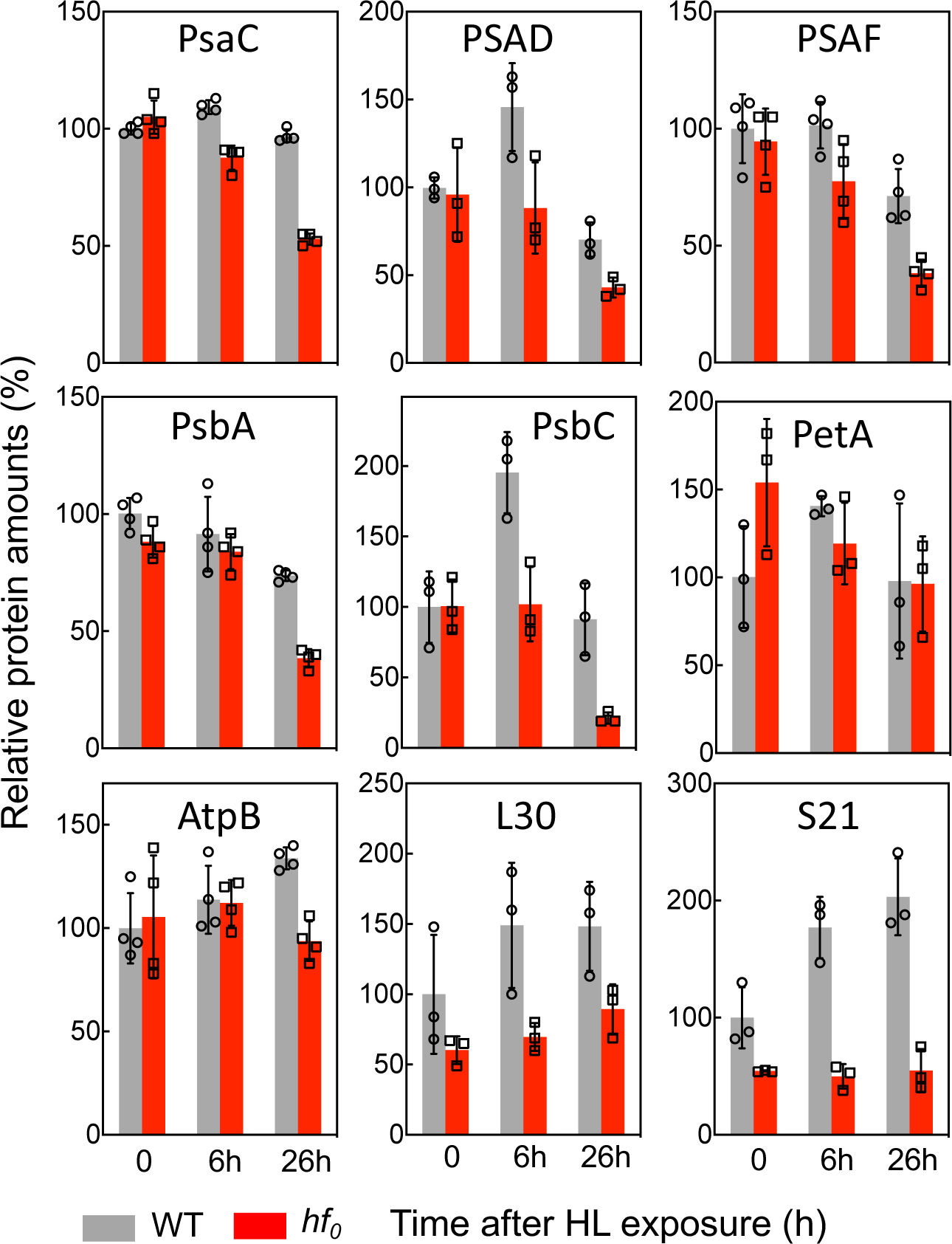
Accumulation of major chloroplast proteins in WT and *hf_0_* upon HL exposure determined by immuno-analysis. Cells were grown photo-autotrophically in photobioreactors in 3% CO_2_-enriched air at LL (7.5 µmol photons m^-2^ s^-1^) and exposed to LL (60 µmol photons m^-2^ s^-1^), for 6h or 26h, respectively corresponding to light intensities of about 20 and 200 µmol photons m^-2^ s^-1^ in flasks (Dang et al., 2014). Protein quantification based on immuno-analysis performed in the linearity range of the antibody detection. Shown are 3 or 4 biological replicates values normalized on the WT protein content measured at t_0_ and means +/-SD.

**Figure 5.**
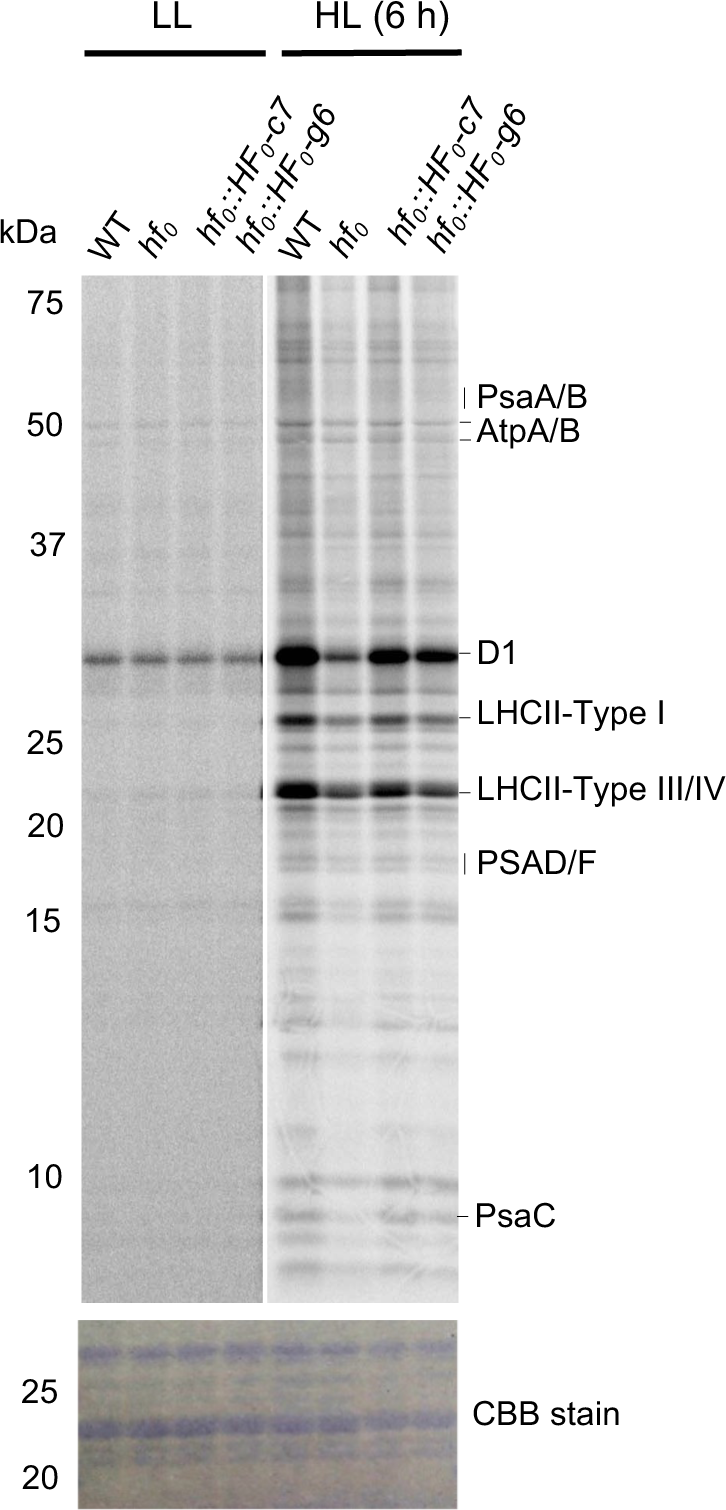
Pulse labeling experiments of thylakoid proteins show a strong decrease of D1 protein synthesis under HL in *hf_0_*. Cells were grown photo-autotrophically at LL up to a cell density of 2. 10^6^ cells mL^-1^ and then switched to HL for 6 h to the cell density of 6. 10^6^ cells mL^-1^, and then labeled for 10 min with ^35^S (Na_2_SO_4_). Upon extraction and SDS-PAGE separation of total cellular proteins, labeled proteins were visualized by autoradiography. Autoradiograms show labeled thylakoid membrane proteins of WT, *hf_0_* and two complemented lines, *hf_0_::HF_0_-*c7 (complemented line obtained by chloroplast transformation) and *hf_0_::HF_0_-*g6 (complemented line obtained by nuclear transformation). The gel was stained with Coomassie brilliant blue before exposing to the imaging plate.

In order to confirm that PSII inhibition observed in *hf_0_*under HL results from a defect in protein synthesis capacity, we used chloramphenicol, an inhibitor of chloroplast protein synthesis. In the absence of chloramphenicol, Φ_PSIImax_ and operating PSII yield progressively decreased upon HL exposure in *hf_0_*but not in wild-type or in complemented lines (Fig. 6 A, C). As previously reported (Schuster et al., 1988), PSII was more strongly inhibited by HL in the presence of chloramphenicol, similar inhibitions being observed in *hf_0_*, wild-type and complemented lines after 7h to 24h exposure (Fig. 6 B,D). Since the chloramphenicol treatment of wild-type somehow mimicked the CreRH22 mutation, we conclude that the HL sensitivity of *hf_0_* likely results from a decreased protein synthesis capacity. Since both turnover and accumulation of some PSII and PSI subunits were affected in HL-acclimated *hf_0_* cells, we measured how PSII and PSI activities are impacted (Supplemental Fig. S7). While a 50% reduction in the maximal PSII yield was observed in *hf_0_* under HL, a smaller decrease in the PSII/PSI ratio was measured, consistent with the existence of reduced PSI activity, although to a lesser extent than PSII inhibition.

**Figure 6.**
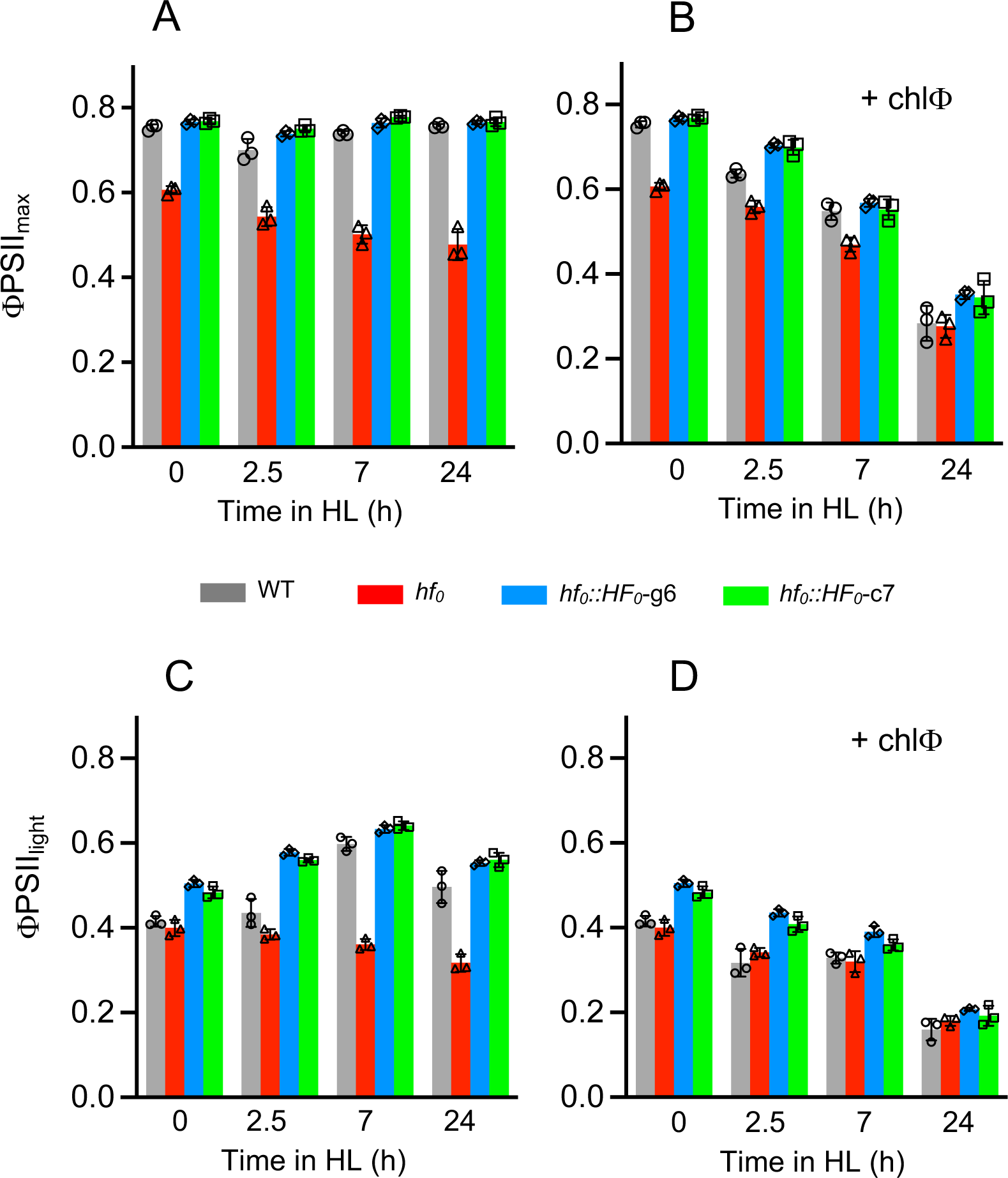
Effect of HL and chloramphenicol addition on photosynthetic activity of the *hf_0_* mutant. Wild-type (WT), *hf_0_* and two complemented lines (*hf_0_::HF_0_-c7* and *hf_0_::HF_0_-g6*) were grown in batch cultures under LL (15 µmol photons m^-2^ s^-1^). At t_0_ cell cultures were switched to HL (240 µmol photons m^-2^ s^-1^) in the absence **(A, C)** or in the presence **(B, D)** of 50 mg. L^-1^ chloramphenicol (chlΦ) added 1h before the HL switch. At different time points after the light switch, maximal PSII yields were measured as the F_v_/F_m_ ratio after 15 min dark adaptation **(A, B)** and PSII operating yields were measured as (F_m’_-F_s_)/F_m’_ following a 3 min light period (430 µmol photons m^-2^ s^-1^). Shown are three biological replicates values and means ± SD.

### Chlorophyll protein complexes are affected in *hf_0_*

In order to gain insight on changes in photosynthesis machinery occurring in *hf_0_* upon HL exposure, we first performed low temperature (77K) chlorophyll fluorescence measurements (Fig. 7). When grown under LL both wild-type and mutant strains showed closely related patterns, with a slightly higher fluorescence emission peak at 712nm (E_712nm_) in the mutant (Fig. 7A). Upon HL exposure, the E_712nm_ peak dramatically increased in *hf_0_* (Fig. 7B). The chlorophyll emission ratio (E_712nm_/E_686nm_) and the F_0_ level (measured as the F_0_/Fm ratio) both strongly increased in the mutant while remaining at nearly constant values in the wild-type (Fig. 7C, D). In order to determine whether the E_712nm_ fluorescence peak increase is related to the phenomenon of state transition, phosphorylation of thylakoid proteins was analyzed (Fig. 7E). During state transitions, reduction of the PQ pool triggers the STT7 kinase, which phosphorylates LHCII and PSII subunits. The phosphorylation status of thylakoid proteins of wild-type and *hf_0_* are similar under LL, and slightly differed under HL (Fig. 7E). However, contrary to what would have been expected if *hf_0_*was more in state 2 than the wild-type, a lower phosphorylation status was observed in the mutant as compared to the wild-type upon HL exposure. Moreover, the blue shift of the 712nm peak observed in HL acclimated *hf_0_* cells (Fig. 7B) is likely due to an increased fluorescence emission from the PSI core or LHCI oligomers, suggesting that part of PSI-LHCI is dissociated into PSI core and LHCI oligomer or that LHCI oligomers accumulate because of the absence of PSI core (Takahashi et al., 2004). We conclude from these experiments that both E_712nm_ and F_0_ increases do not result from a transition to state 2, but likely reflect the presence of LHCI oligomers. We then performed blue native gel electrophoresis of thylakoid protein complexes, and observed no major difference in the distribution of chlorophyll protein complexes in *hf_0_*as compared to the wild-type under LL (Fig. 7F). However, under HL exposure, the pattern of chlorophyll protein complexes of *hf_0_* differed from that of the wild-type, two weak upper bands (PSII-LHCII supercomplexes) disappearing and a lower band (LHCI oligomer) appearing in the mutant, a similar effect being observed when LL wild-type cells were treated with chloramphenicol (Supplemental Fig. S8). Two-dimensional blue native-PAGE showed that a group of polypeptides, associated in wild-type cells to a large complex, are found associated to a smaller complex in *hf_0_* (Fig. 8A, B). Mass spectrometry-based proteomic analysis was performed on a gel strip taken on the 2D gel in the ∼20-35kDa molecular weight range (see red box on Fig. 8B). Various polypeptides were identified, and notably several subunits of PSI (PsaA, PsaB, PsaC, PSAD and PSAF), light-harvesting proteins of PSI (LHCA3, LHCA4, LHCA5, LHCA6, LHCA7), several subunits of PSII (PsbA, PsbB, and PsbC), and one Light Harvesting Complex II (LHCII) polypeptide (LHCBM8), a component of trimeric LHCII (Girolomoni et al., 2017) (Supplemental Data Set S1). Polypeptides with molecular weights comprised between 20 and 35kDa, which corresponds to the gel strip taken for the proteomic analysis, include the five LHCA polypeptides, the two nuclear-encoded PSI subunits (PSAD and PSAF) and LHCBM8. The presence of higher molecular weight polypeptides such as PsaA, PsaB, PsbA, PsbB, PsbD in this area of the gel suggests that some degradation of PSI and PSII photosystems occurred in HL-acclimated *hf_0_* cells. The presence of orphan LHCAs, normally associated with high MW complexes (PSI-LHCI) in a stable manner (Takahashi et al., 2004; Nellaepalli et al., 2021), indicates a defect in PSI organization, in line with 77K chlorophyll fluorescence data (Fig. 7B). It is inferred from this experiment that accumulation of PSI-LHCI and PSII-LHCII supercomplexes, which normally occurs when the synthesis of subunits encoded by nuclear gene and chloroplast genes is properly synchronized, is impaired in *hf_0_*under HL conditions. As a consequence, PSII-LHCII and PSI-LHCI bands disappear, some PSI and PSII reaction centers are degraded and LHCI oligomers accumulate at detectable levels. The increase of LHCII monomer observed in *hf_0_* under HL (Fig. 7F) reflects the fact that in contrast to LHCI, only a small fraction LHCII is normally associated to PSII-LHCII supercomplexes (Tokutsu et al., 2012).

**Figure 7.**
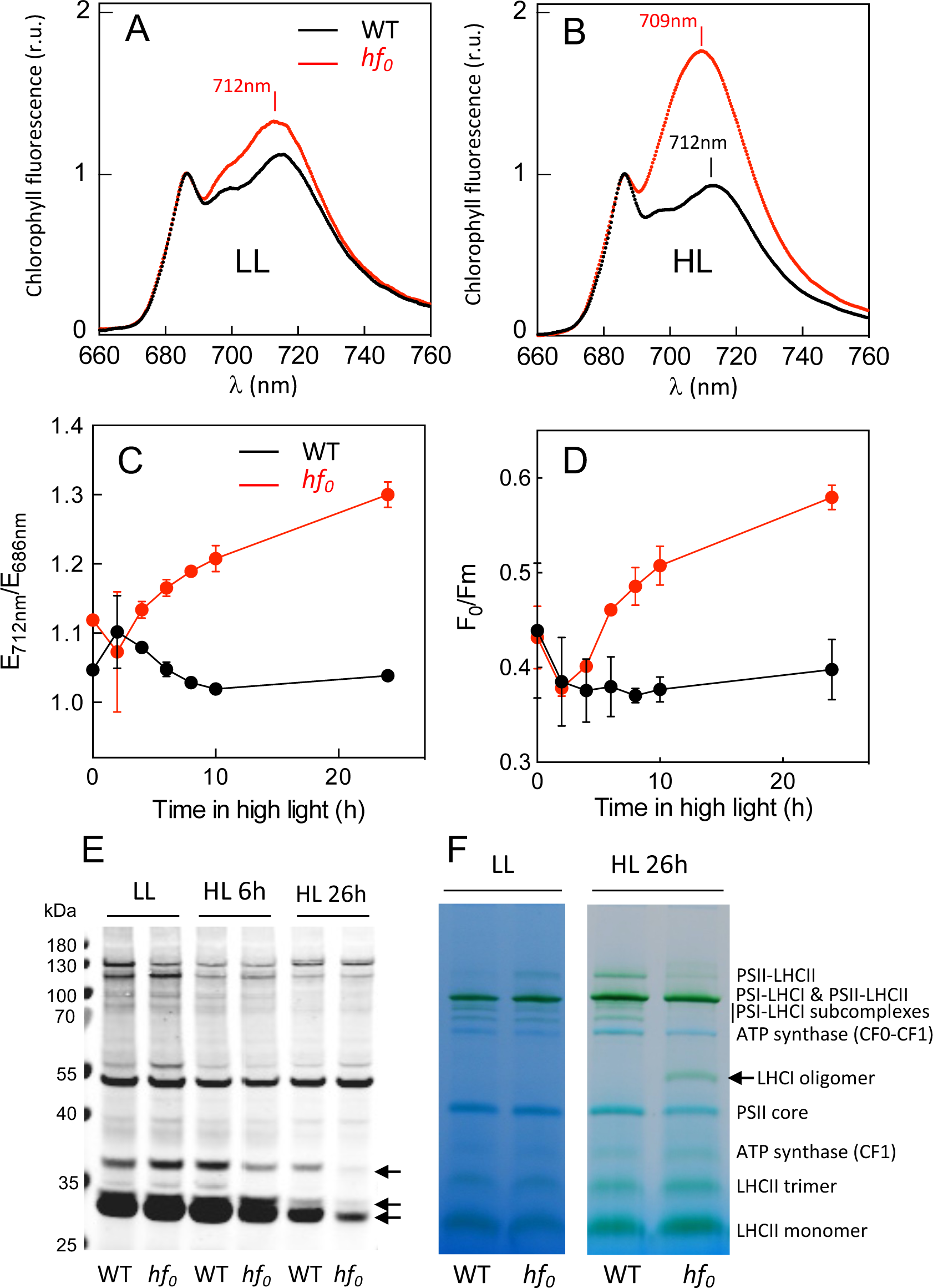
Low temperature chlorophyll fluorescence and chlorophyll-protein complexes are affected in *hf_0_*, but state transitions are not. WT and *hf_0_* cells were grown photo-autotrophically in photobioreactors in 3% CO_2_-enriched air at LL (7.5 µmol photons m^-2^ s^-1^) and then switched to HL (60 µmol photons m^-2^ s^-1^), respectively corresponding to light intensities of about 20 and 200 µmol photons m^-2^ s^-1^ in flasks (Dang et al. 2015). **(A, B)** 77K chlorophyll fluorescence emission spectra of cells grown under LL **(A)** or 26 h after HL switch **(B)**. **(C)** 77K chlorophyll fluorescence emission ratio E_712nm_/E_686nm_, and **(D)** F_0_/F_m_ chlorophyll fluorescence at different time points after HL switch. **(E)** Immunodetection of phosphorylated LHCII using an anti-phospho-threonine antibody. Arrows indicate protein bands with lower abundance in *hf_0_.* (F) Blue-native PAGE of thylakoid complexes from WT and *hf_0_* grown under LL or upon 26h of HL exposure. Two left arrows indicate green bands with lower abundance in *hf_0_*and the right arrow indicates a green band of higher abundance in *hf_0_*.

**Figure 8.**
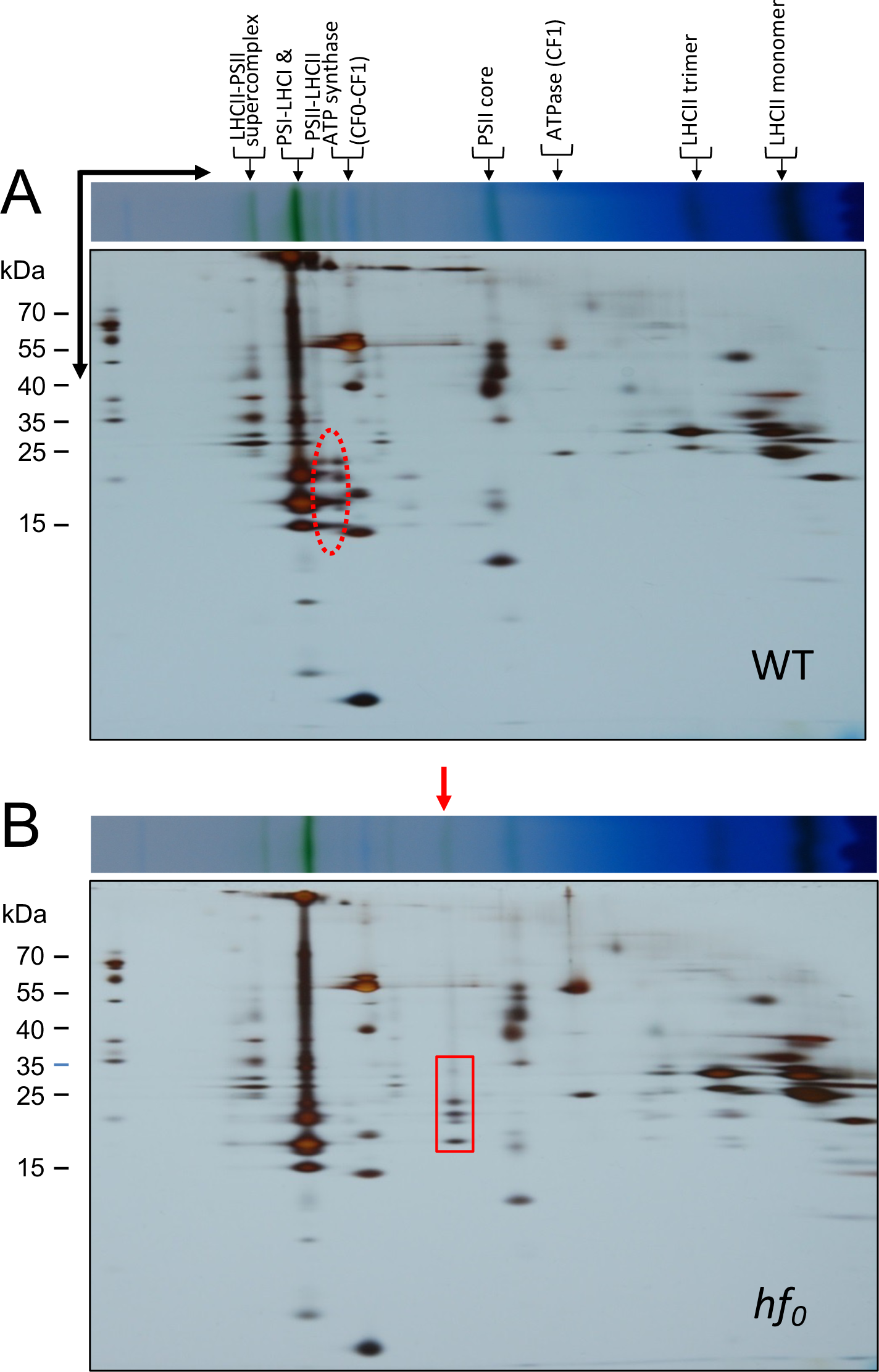
Two-dimensional PAGE of thylakoid proteins from WT and *hf_0_* cells upon HL exposure. WT and *hf_0_* cells were grown photo-autotrophically in batch cultures under LL (40 µmol photons m^-2^ s^-1^) and then exposed to HL (240 µmol photons m^-2^ s^-1^) for 26h. Thylakoid proteins were solubilized with 1% dodecyl-maltoside, separated by blue native PAGE in a first dimension and by SDS-PAGE in a second dimension. The red dotted ellipse highlights proteins associated to high MW PSI-LHCI supercomplexes in the WT **(A)** that migrates as low MW complexes attributed to LHCI oligomers in *hf_0_* (red arrow in **B**). The gel strip in the 20-35kDa MW range shown as a red box **(B)** was taken for proteomic analysis by mass spectrometry (Supplemental Table 1).

### *CreRH22* expression is transiently upregulated during light transients

By searching data related to *CreRH22* (Cre03.g166650) transcript levels in a previous study reporting the dynamics of the *C. reinhardtii* transcriptome in cells synchronized by a night-day cycle (Zones et al., 2015), we found that *CreRH22* expression shows a sharp and transient increase at the beginning of the light period (Supplemental Fig. S9A). In this study a gene cluster named “light stress cluster” characterized by a similar gene expression profile as *CreRH22* was defined. This cluster contains several *HSP* genes (such as *HSP70B* and *HSP90C*), as well as light stress genes such as *PSBS2* or *VTC2* (Zones et al., 2015). Although *CreRH22* was not assigned in this work as a member of the “light stress cluster”, its expression pattern actually meets most criteria used to define this cluster during the light period, the transient expression increase observed during the dark period (reaching 51% of the maximum expression) preventing the classification in this cluster (criteria limit <40%). In order to determine whether *CreRH22* expression is regulated by a light transient independent of circadian regulations, we then performed a dark to light transient on non-synchronized cells, which showed that *CreRH22*, like *HSP70B* and *HSP90C,* is induced by a light (Supplemental Fig. S9B).

### Light-dependent nuclear gene expression is severely impaired in *hf_0_*

RNAseq analysis was then performed in order to determine how the light-induced transcriptional response is affected in the absence of CreRH22. For this purpose, *C. reinhardtii* cells were grown under LL in PBR operated as turbidostats thus allowing acclimation to constant growth conditions, and then switched to HL. Global transcript profiles were analyzed by RNA sequencing after 24h LL acclimation and then after 2h and 24h of HL exposure (Supplemental Data Set S2; Fig. 9). In wild-type cells, 3020 differentially-expressed genes (DEGs) were up-regulated, and 445 down-regulated upon 2h of HL exposure (log 2(Fold Change) > 1 and <-1, respectively) (Fig. 9A). The number of up-regulated DEGs doubled (up to 6023) after 24h under HL, the number of down-regulated DEGs remaining at a low level (576). The effect of HL on gene expression was strongly attenuated in *hf_0_*in comparison to the wild-type, since only 1241 and 1291 genes were up-regulated after 2h and 24h of HL, respectively. In contrast, the number of down-regulated genes doubled as compared to the wild-type. MapMan classification of up-regulated DEGs showed that most represented gene categories are distributed in a similar manner in both strains at different time points (Fig. 9 B,C).

**Figure 9.**
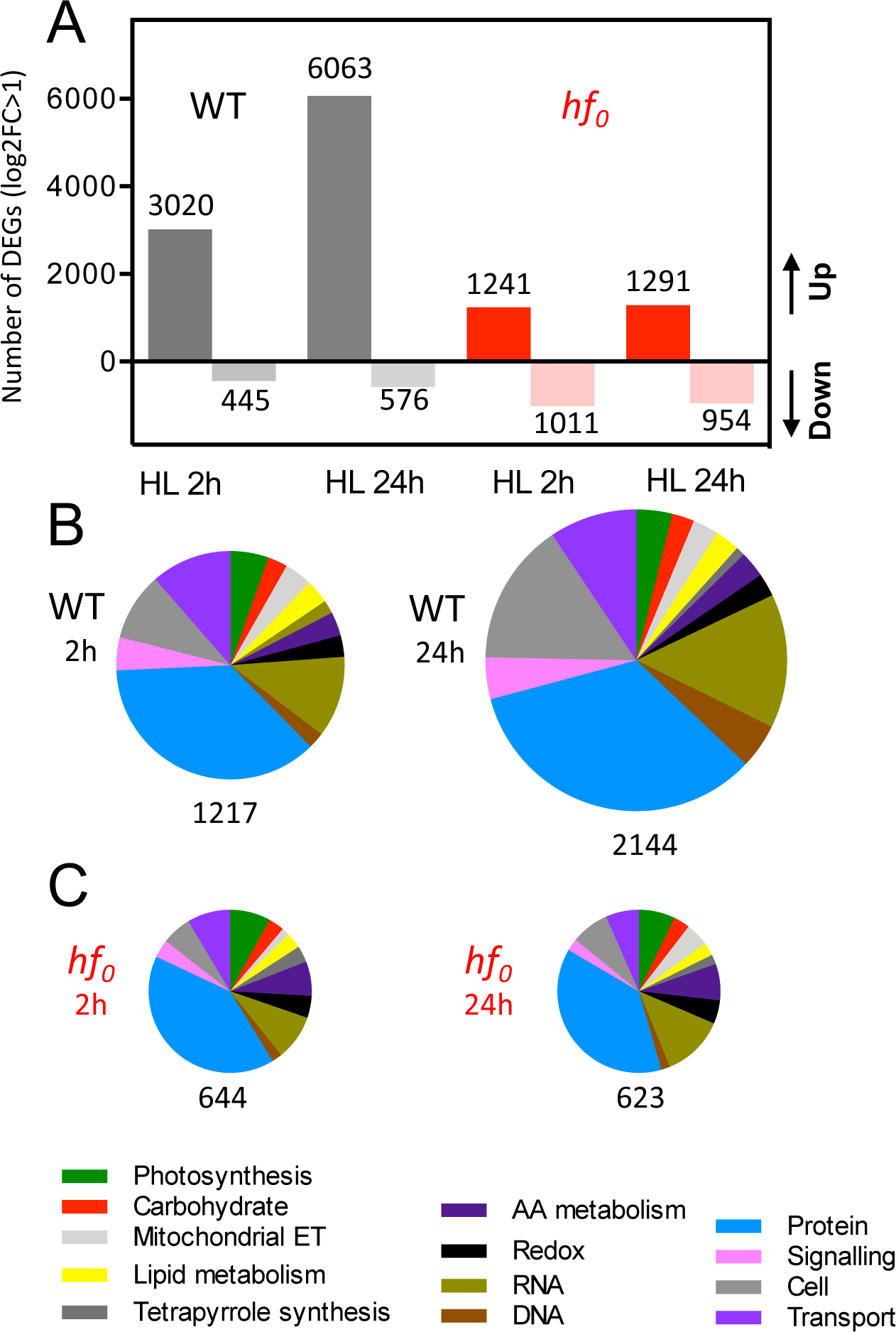
RNAseq analysis of light-induced gene expression in wild-type and *hf_0_ C. reinhardtii* cells. Cells were cultivated in PBRs operated as turbidostats in 3% CO_2_-enriched air under LL (7.5 µmol photons m^-2^ s^-1^) for 24h and then exposed to HL (60 µmol photons m^-2^ s^-1^). Samples for RNAseq analysis were taken from two biological replicates under LL before the HL switch, and the 2h and 24h after the switch. (A) Shown are numbers of up and down Differentially Expressed Genes (DEGs) with a log 2(Fold Change) > 1 and <-1, using a False Discovery Rate (FDR) ≤ 0.001. (B, C) Functional analysis was performed DEGs from WT (A) and *hf_0_* (B) using the MapMan 3.5.1R2 package (Thimm et al., 2004).

Since CreRH22 is involved in chloroplast ribosomal RNA processing, we focused on the expression pattern of ribosomal protein genes (RPGs) of translational machineries located in the cytosol, chloroplast, and mitochondria (Fig. 10). A previous study has established that RPG expression is highly coordinated in the three cellular compartments during light-dark cycles, the expression of cytosolic RPGs peaking at the beginning of the dark period, while the expression of chloroplast RPGS is maximal at the very beginning of the light period (Zones et al., 2015). When wild-type cells were shifted from LL to HL, the maximal RPG expression of cytosolic components was observed under LL and strongly decreased upon HL exposure. In *hf_0_* cells, cytosolic RPG expression was much less affected upon HL exposure (Fig. 10 A,D). The effect of HL on RPG expression of chloroplasts components was dramatically different. Indeed, while expression of chloroplast RPGs was essentially not affected by the HL shift in wild-type cells, a strong and progressive increase in chloroplast RPGs expression was observed in *hf_0_* 2h and 24h after the HL shift (Fig. 10 B,E). On the other hand, mitochondrial RPGs showed a similar expression pattern in the wild-type and *hf_0_* after the HL shift, a similar increase of expression being observed in both strains (Fig. 10 C,F). The dramatic difference between chloroplast RPGs expression observed in *hf_0_*and wild-type cells in response to the HL sharply contrasts with changes in ribosomes abundances, since an increase in ribosomal protein amounts was observed in the wild-type and no change in *hf_0_* (Fig. 4). The increase in chloroplast ribosome abundance observed in wild-type cells in response to HL and the absence of changes in chloroplast RPG expression strongly suggests that chloroplast ribosome abundance is mainly driven by post-transcriptional regulations (*i.e.* increased translation of ribosome subunits). The strong increase in chloroplast RPG expression observed in *hf_0_* upon HL exposure and the absence of changes in ribosome abundance suggests that 23S rRNA processing by CreRH22 is a critical step in the biogenesis of additional ribosomes in wild-type, which cannot be compensated by an increased expression of plastid RPGs in *hf_0_*.

**Figure 10.**
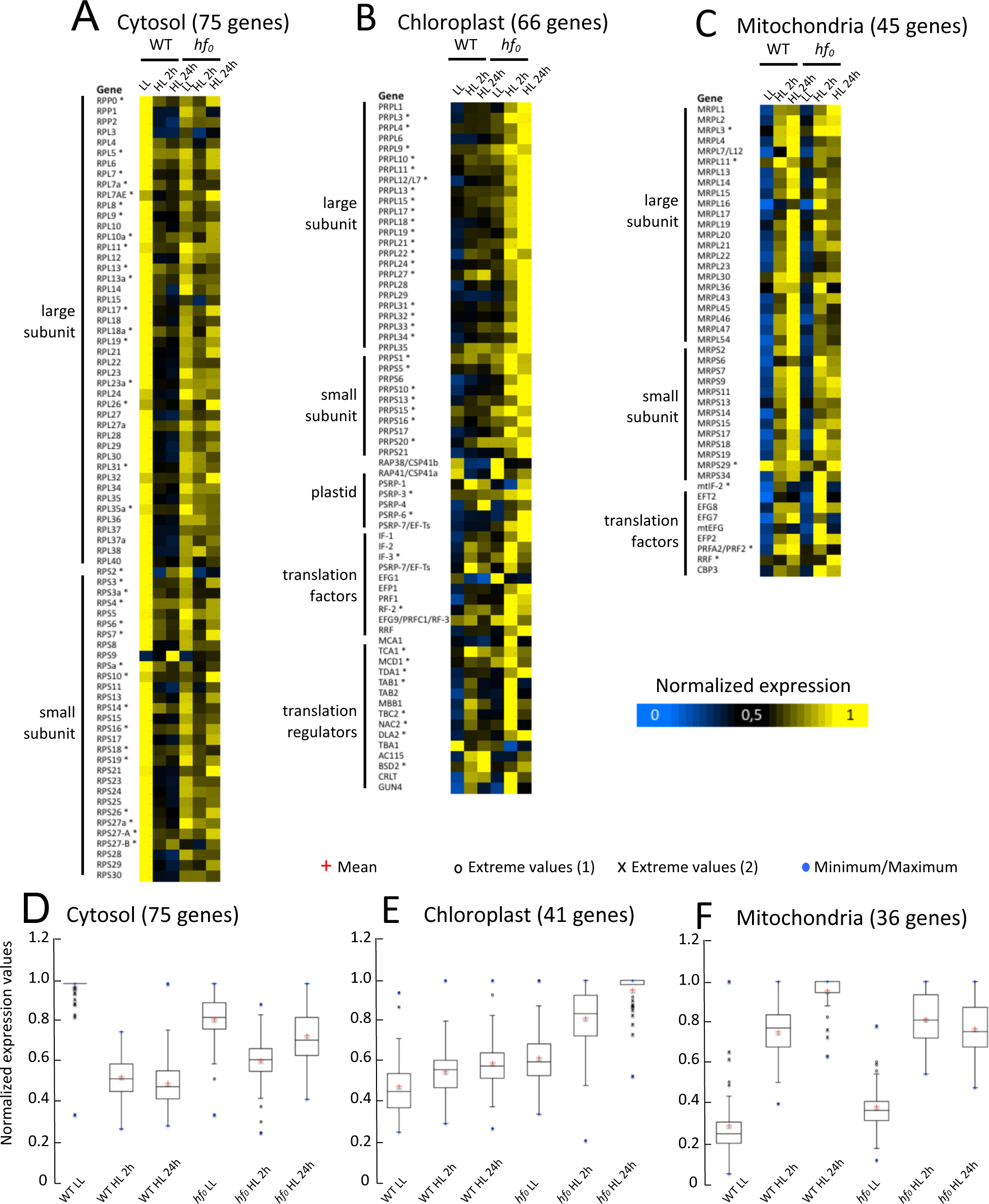
Expression patterns of genes encoding chloroplast-, mitochondria-, and cytosol-targeted ribosomal proteins. **(A to C)** Heat maps depicting relative expression levels of ribosomal protein genes and translational regulators targeted to cytosol **(A)**, chloroplast **(B)**, and mitochondria **(C)**. The maximum expression level for each gene is set to 1. **\*** Genes without significant differential expression (|log2FC|<2 as defined in Methods). **(D to F)** Boxplot of normalized expression values of ribosomal protein genes targeted to cytosol **(D)**, chloroplast **(E)**, and mitochondria **(F)**.

### Biomass productivity is decreased under HL in *hf_0_*

We then aimed to assess the impact of the CreRH22 defect on growth capacity and biomass productivity under different light regimes by performing PBR experiments in a turbidostat mode (Fig. 11). Steady-state dilution rates of cultures maintained at constant biomass concentration allowed to determine specific growth rate, which increased as a function of light intensity in both wild-type and *hf_0_* cell cultures until a saturation plateau, which reached a 40% lower value in *hf_0_* as compared to the wild-type (Fig. 11A). Remarkably, the decrease in the cellular chlorophyll content observed upon HL acclimation of *C. reinhardtii* cells (Bonente et al., 2012) was strongly diminished in *hf_0_* (Fig. 11B) indicating a strong impairment of light acclimation in the mutant. Under stress conditions, such as nutrient depletion (Siaut et al., 2011) or HL (Goold et al., 2016), *Chlamydomonas* cells accumulate starch and oil, which is generally viewed as the result of an imbalance between photosynthesis and growth capacities. While intracellular starch and oil reached high levels in the wild-type at HL, starch accumulation was much lower in *hf_0_* (Fig. 11D), the difference in oil contents being less marked (Fig. 11E). We conclude from this experiment that CreRH22 is critical for Chlamydomonas to acclimate to HL conditions and achieve maximal biomass productivity.

**Figure 11.**
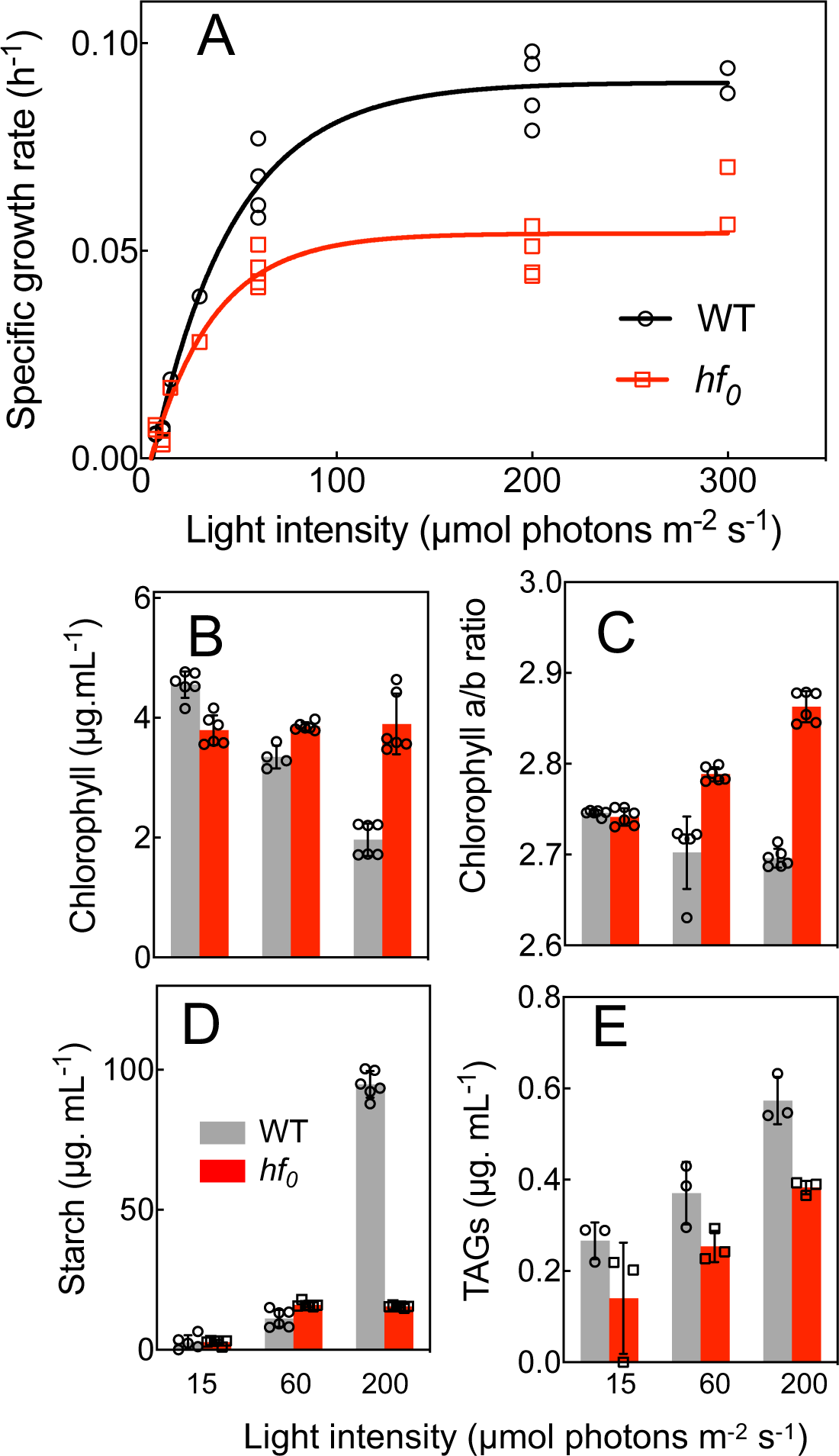
Biomass productivity and concentrations of intracellular starch and TAGs are reduced in *hf_0_*. Wild-type and *hf_0_* cells were grown in PBRs operated as turbidostats in 3% CO_2_-enriched air and maintained at a constant biomass concentration (≈ 1.5 x 10^6^ cells mL^-1^) by continuous addition of fresh medium. **(A)** Specific growth rates were determined from dilution rates measured at different light intensities (ranging from 7.5 up to 300 µmol photons m^-2^ s^-1^) following at least 24h stabilization. **(B)** Chlorophyll content, **(C)** chlorophyll a/b ratio, **(D)** intracellular starch, and **(E)** intracellular TAG contents were determined on cell samples following at least 24h stabilization at the specified light intensity (shown are means ± SD, 3<n<6).

## DISCUSSION

We have shown here that CreRH22, a DEAD-box RNA helicase targeted to the chloroplast and involved in ribosomal RNA processing and ribosome biogenesis, has a critical function during HL acclimation in the green unicellular alga *C. reinhardtii*. CreRH22 belongs to the GreenCut2 (Karpowicz et al., 2011) and shows similar gene expression profile as a transiently expressed cluster of light stress genes (Zones et al., 2015). We propose that rapid induction of CreRH22 upon HL exposure promotes ribosome assembly, thus increasing the translation capacity and allowing synthesis of plastid-encoded proteins of the photosynthesis machinery, and particularly the D1 protein of PSII to avoid photoinhibition. In the absence of CreRH22, plastid-encoded PSII subunits (and to a lesser extent the PSI subunits) are not synthesized at a sufficient rate during HL exposure, thus leading to an imbalance between the synthesis of nuclear- and plastid-encoded subunits of photosystems, to the accumulation of lower levels of PSII-LHCII and PSI-LHCI complexes and to the accumulation of oligomeric LHCI, thus explaining the increase in the F_0_ chlorophyll fluorescence. As a consequence, light-induced gene expression, photosynthesis and biomass production are strongly affected under HL (Figs. 11 and 12).

**Figure 12.**
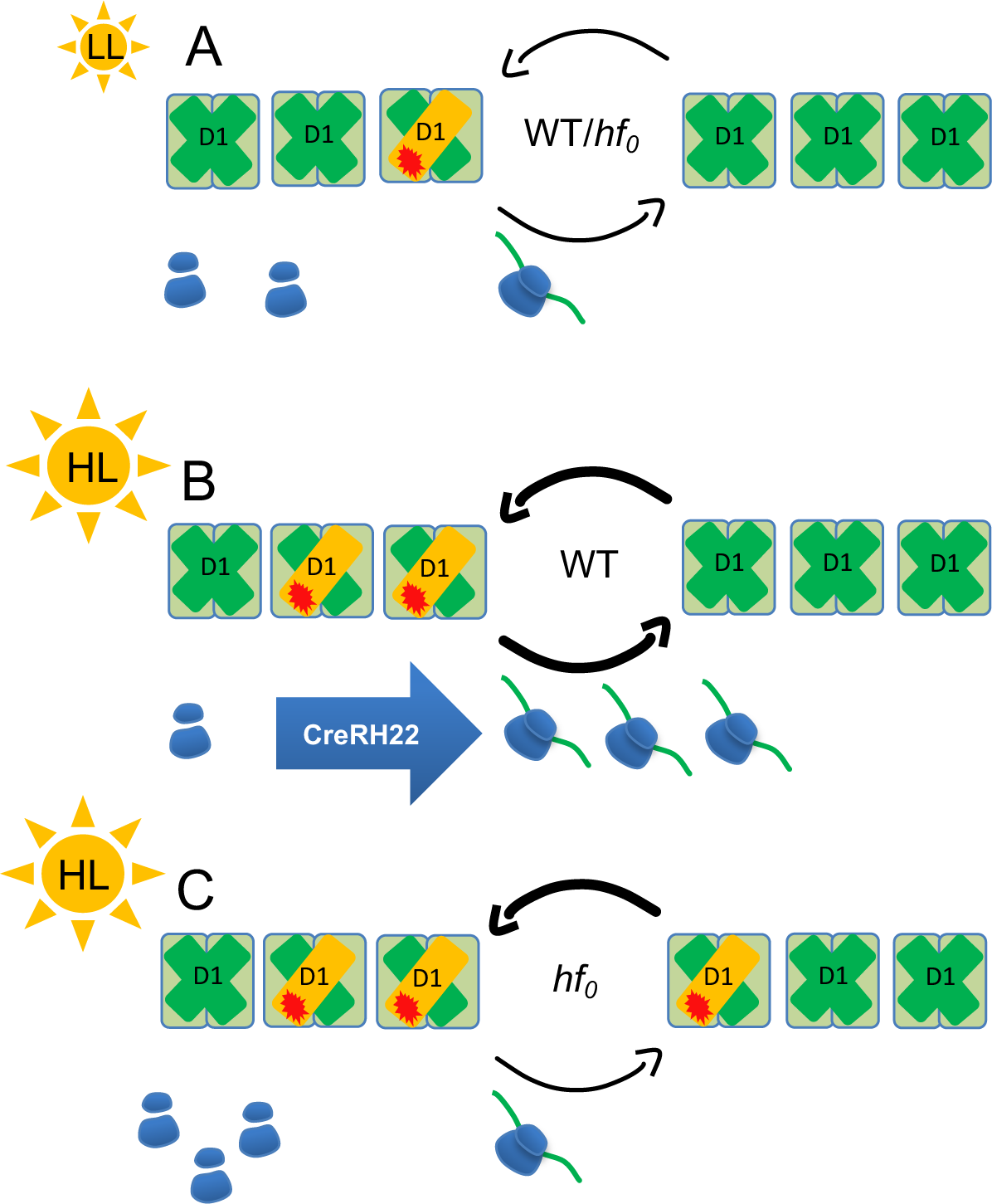
Hypothetical scheme proposed to explain the *hf_0_* mutant phenotype. The *hf_0_* mutant is defected in a plastidial DEAD-box RNA helicase (CreRH22) involved in the processing of plastidial rRNA operon and ribosome biogenesis. **(A)** Under LL the turnover of chloroplast proteins is low and the translation capacity is sufficient to maintain photosynthesis at similar levels in the wild-type (WT) and *hf_0_*. **(B, C)** Under HL exposure, the amount of chloroplast ribosomes increases in the WT but remains constant in the mutant. In these conditions, a much higher rate of protein synthesis is needed due to the permanent degradation and synthesis of the D1 protein and to the biosynthesis of additional PSII subunits and to a lesser extent PSI subunits. In these conditions, the *hf_0_*mutant is unable to synthesize photosynthetic proteins at a sufficient rate, thus resulting in an inhibition of photosynthesis.

### Physiological functions of plastidial DEAD-box RNA helicases

DEAD-box RNA helicases are involved in different aspects of RNA metabolism and participate to many cellular functions (Linder and Jankowsky, 2011). In Arabidopsis, among 58 DEAD-box RNA helicases identified in the genome, ten and eight are predicted to be respectively targeted to chloroplasts or mitochondria and involved in plant response to abiotic stresses (Nawaz and Kang, 2017). Proteomic analysis revealed the presence of six of the plastidial DEAD-box RNA helicases, including RH22, in high molecular mass stromal complexes, thus suggesting an involvement of these proteins in ribosome biogenesis (Olinares et al., 2010), which was confirmed for RH3 and RH22 by studies of Arabidopsis mutants (Asakura et al., 2012; Chi et al., 2012). However, despite the presence of DEAD-box RNA helicases in the chloroplast stroma, a possible function of these proteins in regulating photosynthesis has not been documented so far. Indeed, loss of the Arabidopsis *RH22* gene is lethal, and knock-down mutants show a severe growth phenotype (Chi et al., 2012). In contrast, disruption of the *C. reinhardtii* gene confers a light-dependent phenotype, the mutant growing normally under LL and at a slower rate than the wild-type under moderate to HL. Such a phenotype difference between plants and unicellular algae may be related to the requirement of a high protein synthesis capacity during the process of plastid biogenesis and/or differentiation occurring during plant greening (Sun and Zerges, 2015). In this context, any disturbance in the translation machinery has a strong impact on plant chloroplast biogenesis. However, unicellular organisms such as microalgae do not experience such a greening process since chloroplasts divide as green organelles (Jarvis and Lopez-Juez, 2013). Both CreRH22 and AtRH22 are thus likely needed when active protein synthesis is required, either during chloroplast biogenesis in plants, or during the PSII repair cycle in HL conditions in algae and most probably also in plants. However, the later function is difficult to assess due to the strong phenotype of the Arabidopsis RH22 mutant.

### CreRH22 is part of a network of proteins involved in ribosomal biogenesis

Remarkably, five out of the seven DEAD-box RNA helicases predicted to be plastid targeted in Chlamydomonas, including CReRH22, belong to the gene cluster 1 (Supplemental Fig. S3), defined as genes showing a strong induction during the first hour of illumination followed by a sharp decrease (Zones et al., 2015). These genes are expressed before the maximal expression of most ribosome protein genes, which peaked at two hours of light exposure (Zones et al., 2015), in line with a function in ribosome biogenesis. Very recently, four of these DEAD-box RNA helicases, including CreRH22, were identified as interacting with chloroplast ribosome proteins (Westrich et al., 2021) (Supplemental Fig. S3). Chloroplast ribosomes are made of a small subunit (30S) containing the 16S ribosomal RNA and a large subunit (50S) containing 23S ribosomal RNA. While AtRH22 in Arabidopsis (Chi et al., 2012) and CreRH22 in Chlamydomonas (this work) are involved in 23S ribosomal RNA maturation, a bacterial-type ribosome binding factor (RbfA) was proposed in Arabidopsis to participate in the processing of 16S ribosomal RNA. RbfA also belongs to the GreenCut, and was shown in Arabidopsis to be essential for chloroplast development and photoautotrophic growth (Fristedt et al., 2014). As it is the case for CreRH22, the Chlamydomonas RBF1 homolog (Cre02.g145000) belongs to the gene cluster 1 (Zones et al., 2015), was found in high molecular mass stromal complexes (Olinares et al., 2010), and was enriched in 30S ribosomal fractions interactome (Westrich et al., 2021). Together, these studies suggest that chloroplast CreRH22, in association with other plastid-targeted DEAD-box RNA helicases, is involved in ribosome biogenesis in response to the light supply. CreRH22 would be part of a complex network of regulatory proteins including other plastid targeted DEAD-box RNA helicases and other factors such as RBF1, involved in a coordinated increase in the translation machinery capacity required when algal cells are exposed to HL.

### Crosstalk between transcriptional and post-transcriptional regulations during HL acclimation

*Chlamydomonas* implements diverse regulation strategies and photoprotection mechanisms to avoid photo-oxidative damages. PSII photoinhibition occurs when the rate of PSII damage exceeds the rate of repair. The PSII damage and repair cycle is actually part of a dynamic machinery, acting on a principle similar to that of a circuit breaker, which in coordination with other regulatory processes operating on shorter time scales (such as photosynthetic control or qE), protects PSI from irreversible photo-damages (Krieger-Liszkay et al., 2000; Suorsa et al., 2012; Tikkanen et al., 2014; Chaux et al., 2015). Optimal functioning of photosynthesis under HL therefore relies on a fine poise between PSII degradation and synthesis, the latter depending on the translation capacity, which in bacterial systems greatly relies on the level of free ribosomes (Kim et al., 2020). In such a scenario, chloroplasts need to improve the translation capacity to increase the production of photosystem subunits, particularly D1, in response to HL to avoid severe photo-oxidative damages. CreRH22, together with other plastid-targeted DEAD-box RNA helicases and other factors such as RBF1, may contribute to finely tuning the translation capacity to the protein synthesis demand. Ribosomes represent important energy cost and resource allocation for cells, and in bacteria, the number of ribosomes is the result of a trade-off in the use of nutrients between protein synthesis capacity and production of metabolic enzymes (Kim et al., 2020). In bacteria and yeasts, allocation of resources towards ribosome synthesis is tuned to optimize growth (Chure and Cremer, 2022). Photoautotrophic organisms are likely to have similar constrains, especially when it comes to tune ribosomal content devoted to photosynthetic proteins when light is the sole source of energy given the strong turnover rate of photosynthetic proteins (Li et al., 2018). Optimizing the number of chloroplast ribosomes to available light resources is most likely to be critical to maintain photosynthesis functioning and generate the required energy to optimize fitness during adverse conditions of light fluctuations, and the DEAD-box RNA helicase CreRH22 appears to play a prominent role in such a regulation (Fig. 12).

Chloroplasts are semi-autonomous organelles of prokaryotic origin housing photosynthesis in plant and algal cells. Most of the genes initially present in the endosymbiotic ancestor of chloroplasts have been relocated to the nuclear genome, leading to a situation in which multi-subunit photosynthetic complexes and ribosomes are partly encoded by nuclear and plastid genes. Therefore, a coordinated control of gene expression and of protein synthesis between these two compartments is needed to ensure a well-orchestrated assembly of these complexes and an optimal functioning of photosynthesis (Stern et al., 2010). The expression of plastid genes is tightly controlled by complex regulatory mechanisms involving more than hundred nucleus-encoded proteins (Stern et al., 2010), and plastid-targeted DEAD-box RNA helicase most likely contribute to such a control.

An intriguing question relates to the origin of the strong changes in transcriptional regulation that occur during light acclimation in *hf*_0_. The expression of nuclear genes depends on a retrograde signaling originating from the chloroplast, the origin and nature of which being a matter of intense debate. It has been alternatively proposed that the redox state of the PQ pool or the redox state of PSI acceptor side may contribute (Escoubas et al., 1995; Allen and Pfannschmidt, 2000; Pfalz et al., 2012; Dietzel et al., 2015). In Arabidopsis leaves, the redox state of compounds on the reducing side of PSI was considered to be more important than PQ-mediated redox signals in controlling the expression of nuclear genes (Piippo et al., 2006). The redox status of the PQ pool can be assessed from chlorophyll fluorescence measurements in certain conditions through the use of the parameter 1-qL (Kramer et al., 2004). However, the high chlorophyll fluorescence of the *hf_0_* mutant, which results from the presence of non-connected antennae, is not consistent with the “lake model” hypothesis that considers fully connected units (Kramer et al., 2004), thus preventing the use of this parameter in this context. Since phosphorylation of LHCII antennae, which depends on the reduction state of the PQ pool (Depege et al., 2003), is not increased in *hf_0_* (Fig. 7. E), it seems likely that the redox state of the PQ pool is not increased in *hf_0_*. In fact, a lower redox state of the PQ pool is likely to occur in *hf_0_*since PSII is more strongly inhibited than PSI under HL (Supplemental Fig. S7B). In this context, we propose that the strong impairment of the transcriptome response observed in *hf_0_* under HL would be rather due to limited over-reduction of PSI acceptors under HL resulting from a lower synthesis of photosystems.

Among the 32 Chlamydomonas DEAD-box RNA helicases listed in the Supplemental Fig. S3, 11 belong to a gene cluster (cluster 1) containing genes characterized by a strong induction during the first hour of illumination followed by a sharp decrease (Zones et al., 2015), and four out of these 11 gene products were recently identified as physically interacting with plastid ribosomes (Westrich et al., 2021). It seems therefore likely that, in addition of CreRH22, several other plastid-targeted DEAD-box RNA helicases may participate, via the nuclear control of the plastid protein synthesis, to the coordination between nuclear and plastid compartments, which is critical for a correct assemblage of chloroplast complexes, particularly in changing light conditions. Recently, many DEAD-box RNA helicases have been identified and partially characterized in crop plant genomes, including rice (Lu et al., 2020), wheat (Ru et al., 2021), *Medicago truncatula* (Cheng et al., 2021) and *Brassica rapa* (Nawaz et al., 2021), and plastid-targeted DEAD-box RNA helicases have been proposed to play a role in the response to abiotic stress such a drought, salinity or cold (Nawaz and Kang, 2017). As light is as a key parameter that potentiates the effects of abiotic stress on photosynthesis (Schwenkert, in press), plastid-targeted DEAD-box RNA helicases may represent promising targets towards improving the productivity of crops in stress conditions.

## MATERIALS AND METHODS

### Strains and culture conditions

*C. reinhardtii* strains (CC124 called wild-type, the *hf_0_*mutant and complemented strains) were grown photo-autotrophically on a minimal medium (Harris, 2009). Batch cultures were grown on a rotary shaker in Erlenmeyer flasks (100 mL) placed in a thermo-regulated (25°C) incubator (Multitron, Infors) under continuous illumination (40 or 240 µmol photons m^-2^ s^-1^) in the presence of 2%CO_2_ enriched air. For continuous photoautotrophic growth experiment, cells were cultivated in four autoclavable 1L PBRs (BIOSTAT® Aplus, Sartorius Stedim Biotech) equipped with a biomass probe (Excell probe, Exner, measuring O.D at 880 nm with a 2 cm light path) and operated as turbidostats (Dang et al., 2014). Light was supplied by eight fluorescent tubes (Osram Dulux L 18W) disposed radially around the PBR to reach light intensities (measured at the surface of the PBR). Growth rates (d^-1^) were calculated by dividing the daily dilution volume by the PBR volume (1L) and biomass productivity from dry weight measurements.

### Mutant screening, isolation and molecular characterization

The *C. reinhardtii* strain CC124 (*mt^-^ nit1 nit2*) was used as a genetic background to generate an insertion mutant library as described previously (Tolleter et al., 2011). The pSL-X plasmid harboring the paromomycin resistance cassette *AphVIII* was linearized by *KpnI* and used for nuclear transformation by electroporation. Around 12,000 random insertion mutants were isolated on selective medium and mutant phenotypes were analyzed by chlorophyll fluorescence using a homemade imaging device. The *hf_0_* mutants showed a high F_0_ chlorophyll fluorescence level. The insertion site of the paromomycin cassette was identified on genomic DNA upon phenol/chloroform extraction (Tolleter et al., 2011). First, Genome walker protocol was adapted as follows from Universal GenomeWalker^TM^ kit (Clontech). 250ng of gDNA were digested overnight by 80U of *Afe*I (New England Biolabs). The restriction enzyme was eliminated by precipitation with 1/10 (v/v) of ammonium acetate 2.5M (pH 5.6) and 1V of cold EtOH. Digested DNA was dephosphorylated for 1h by 10U of antartic phosphatase (New England Biolabs). DNA adaptator was ligated overnight with 2,000 u of T4 DNA ligase (New England Biolabs). Two PCRs were carried out by using nested primers designed on the adaptator and on the paromomycin resistance gene using the Advantage® GC genomic LA polymerase mix (Clontech). PCR products were subcloned for sequencing and blasted on the *C. reinhardtii* genome, thus leading to the identification of the Cre03.g166650 gene (thereafter called *CreRH22*).

### Southern blot analysis

Genomic DNA (8_μ_g) was digested by *NcoI* (New England Biolabs) overnight and loaded on 0.8% agarose gel. After migration, gel was depurinated in 0.25M HCl for 10 min, denatured in 0.5M NaOH/1.5M NaCl, and neutralizated for 30 min in 0.5M Tris-HCl (pH 7.5), 1.5M NaCl. The DNA was transferred overnight onto a nylon membrane using a 10 x SSC buffer, and was fixed onto the membrane by a UV treatment (1,200W, UV Crosslinker, UVP Laboratory Product, UK). Paromomycin probe was labeled with dUTP-DIG during PCR reaction (PCR DIG probe synthesis kit, Roche; see table primers list). Membrane was pre-hybridized 1h at 50°C in a DIG eazy Hyb (Roche) and then hybridized overnight at 50°C in same buffer using a denatured probe. Two washing steps were realized, first using a 0.2 x SSC buffer containing 0.1% SDS (5 min at RT), and then a 1 x SSC buffer containing 0.1% SDS (20 min at 65°C). Detection was done by Western Blot with anti-digoxigenin antibody, coupled with alkaline phosphatase. We followed Roche procedure: 5 min in washing buffer, 30 min in blocking buffer, hybridization during 1h (1/10e4 anti-DIG in blocking solution), two washing step of 15 min. Membrane was then equilibrated with detection buffer (pH 9.5) before to add chemiluminescent substrate CSPD® (disodium 3-(4-methoxyspiro {1,2-dioxetane-3,2’-(5’-chloro)tricyclo [3.3.1.1^3,7^]decan}-4-yl) phenyl phosphate) (Roche). The chemio-luminescent signal was detected by using a G-Box (Syngene).

### Mutant complementation

Complementation of the *hf_0_* mutant was first attempted by nuclear transformation by cloning the genomic *CreRH22.* The CreRH22 gene was amplified by Phusion® High-Fidelity PCR Master Mix with CDS helicase primers (Supplemental Fig. S10), subcloned into the topoXL vector, and transferred into the pSL vector carrying the hygromycin resistance cassette using *EcoRV/SpeI* sites (Berthold et al., 2002), and the CreRH22 gene under the control of *psa*D promoter and terminator. The vector was linearized by *KpnI* and transformation was performed as previously described (Tolleter et al., 2011). Transformants were selected on hygromycin (7.5µg mL^-1^) and further analyzed for chlorophyll fluorescence properties. Complementation was also attempted by means of chloroplast transformation. A synthetic *CreRH22* gene optimized for the *Chlamydomonas* chloroplast codon usage (Supplemental Fig. S5) was cloned into the pLM21 vector under the control of the *psa*A promoter using *BglII* restriction sites (Michelet et al., 2011; Tibiletti et al., 2016). Cells were grown at 25°C under continuous light (100 µmol photons m² s^-1^) in liquid Tris-Acetate-Phosphate (TAP) medium until a density of 4.10^6^ cells mL^-1^ and then spread onto agar plates. Bombardment was realized at 7 bars with 10µg DNA of the PLM21 vector bound to gold carrier particles (Seashell Technology kit). Transformants were selected on a medium containing 100 µg mL^-1^ spectinomycin, and replated on increasing spectinomycin concentrations (up to 500 µg mL^-1^) until homoplasmy was reached, as controlled by PCR (Baltz et al., 2014).

### RNA extraction and reverse transcription

After harvesting cells by centrifugation, cell pellets were frozen in liquid nitrogen and resuspended in 0.5 mL of RNA reagent (Life Technologies™) to isolate total RNAs. Cells were broken (two times for 10s at 5500 rpm, 4°C) using ceramic beads in a 2 mL tube using a Precellys24® homogenizer (Bertin technologies, France). 100μL of NaCl 3M and 300 μL of chloroform were added sequentially. After centrifugation (13,000g, 4°C) the upper aqueous phase was mixed with isopropanol in order to precipitate nucleic acids. After washing with 70% EtOH RNA samples were air dried. A DNAse treatment was done with TURBO® DNAse (Ambion®) followed by purification with NucleoSpin® RNA clean-up (Macherey Nagel). Reverse transcription was performed on 1µg of the total RNA using the SuperScript® vilo™ cDNA synthesis kit (Life Technologies™). Expression of *CreRH22* was monitored by PCR on cDNA using TaKaRa LA Taq® DNA Polymerase. *CBLP2* (RACK1 Cre06.g278222.t1.1) was used as a control (Supplemental Fig. S10).

### Northern blot and real time quantitative PCR

For Norther blots, 5µg of total RNA were separated on 1.4% agarose denatured gel (2% formaldehyde), transferred on nylon membrane and hybridized with DIG-labelled probes obtained by PCR DIG Probe Synthesis Kit (Roche) with specific primers (see table primers list). The membrane was pre-hybridized and hybridized at 50°C in DIG eazy Hyb (Roche). Two double washing steps were realized using 0.1% SDS, 0.2x SSC, during 5 min at room temperature and 20 min at 65°C. Detection was performed using an anti-digoxigenin antibody, coupled with alkaline phosphatase. RT-qPCR reactions were realized on LightCycler® 480 (Roche) with MESA FAST qPCR MasterMix Plus for SYBR® Assay No ROX (Eurogentec). The cycling conditions used were as follows: 95°C for 10 min, then 45 cycles at 95°C for 10 s, 60°C for 15 s, and 72°C for 10 s. The relative transcript ratio were calculated based on the 2^-ΔΔCT^ method (Livak and Schmittgen, 2001), with average cycle threshold obtained based on triplicate measurements. Ribosomal protein gene *rpl36* was used as chloroplast reference gene.

### RNA-seq experiments

10 mL of PBR turbidostat cultures of both wild-type and *hf_0_*(with a cell density of 3 x 10^6^ cells mL^-1^) were harvested and centrifuged (4000 rpm, 1 min, 4°C). Samples of two replicate cultures (two independent PBR experiments) were taken after 24 h cultivation under 30 µmol photons m^-2^ s^-1^ and then 2h and 24h after a switch to 240 µmol photons m^-2^ s^-1^. Total RNAs of all samples (12 samples in total) were extracted as described above. Each library was sequenced using an Illumina Sequencer generating 23.3-25 10^6^ 50 nt single end reads for each sample. The sequencing reactions and RNA-Seq data analysis were performed at Beijing Genome Institute, Hong Kong. For transcriptomic analysis, reads were aligned onto the *C. reinhardtii* genome assembly version v5 using SOAPaligner/SOAP2 (Li et al., 2008; Li et al., 2009). No more than two mismatches were allowed in the alignment. For each sample 17.2-19.8 10^6^ reads were mapped to single copy sequences on the *C. reinhardtii* genome. NoiSeq (Tarazona et al., 2011) was applied to screen Differentially Expressed Genes (DEGs) between two groups. We used a False Discovery Rate (FDR) ≤ 0.001 and the absolute value of ≥ 1 as the threshold to judge the significance of gene expression difference. These DEGs were submitted to functional analysis using the MapMan 3.5.1R2 (Thimm et al., 2004) software package to attribute DEGs to biological pathways.

### Starch and oil measurements

Starch extraction was performed using the method described by Klein and Betz (1978). One ml of culture was centrifuged (14,000 rpm, 2 min), re-suspended in 1ml of methanol and centrifuged again. Chlorophyll concentration was estimated directly on methanol supernatants by absorbance at 663 and 645 nm. Pellets were heated for 15 min at 120°C for starch solubilisation after the addition of 400µL water. 200µL of amyloglucosidase solution Starch Assay Reagent (Sigma-Aldrich, Saint-Louis, MO, USA) were then added and incubated at 55°C for 1h. Glucose was subsequently assayed using an automated sugar analyser (Ysi model 2700 select, Yellow springs, OH, USA). Lipid extraction and TAG analysis were performed as previously described (Siaut et al., 2011).

### Immunoblot analysis

Upon harvesting, 20x10^6^ cells were harvested and frozen from PBR cultures. Totals proteins were extracted with SDS 0.2% and acetone 80%. The pellets were treated with a denaturing buffer from Thermofisher (LDS Sample Buffer supplied with DTT) and heated during 20 min at 70°C. Gel electrophoresis was performed using a NuPAGE Novex System (10*%* Bis*-*Tris gel, MES running buffer) and proteins transfer using iBlot® 2 Dry Blotting System. Depending on the sensitivity of the antibody, detection was based on chemiluminescence using HRP conjugated antibody, chemiluminescent substrate and a GBOX imaging system (Syngene), or fluorescent detection using a Fluorescent Alexa Fluor 680 dye conjugated antibody and Odyssey imaging system (LiCor). Antibodies against PsbA, PsbC, PsaC, PSAD, AtpB, and PetA were purchased from Agrisera and the phosphothreonin antibody was purchased from InvitroGen. The PSAF antibodies was kindly supplied by Michel Goldschmidt-Clermont (University of Geneva, Switzerland), and L30 and S21 antibodies by William Zerges (Concordia University, Montreal, Canada). Immuno-quantification was performed in the linear range of each antibody response. Protein amounts loaded for immunoblots were 8µg (PsaC, PSAD, PSAF, PsbC), 4µg (PetA, L30), and 1.6µg (AtpB, PsbA). Densitometries were analyzed by using the GeneTools analysis software (Syngene) for chemiluminescence measurements or the Odyssey analysis software (LiCor) for fluorescence measurements. Quantifications were based on biological triplicates and expressed as relative values of the average wild-type protein level at LL.

### Blue native PAGE and two-dimensional analysis of thylakoid proteins

Thylakoid membranes were purified under dim light at 4°C by using a protocol adapted from (Chua and Bennoun, 1975). Cells were harvested during exponential growth (about 5 10^7^ cells) by centrifugation (5,000 rpm, 5 min, 4°C). Cell pellets were resuspended in 5mL of buffer A containing 5 mM HEPES-KOH (pH 7.5), 0.3M sucrose, 10 mM EDTA, Protease Inhibitor Cocktail (SigmaAldrich). Cells were disrupted using a French Press (Aminco) and centrifuged (12,000g, 10 min, 4°C). Cell pellets were resuspended in buffer A containing 1.8 M sucrose, and blended with a potter homogenizer. The homogenate was covered with 4 mL of buffer A without EDTA and containing 1.3M sucrose and 4 mL of buffer A containing 0.5 M sucrose. Upon ultracentrifugation (274,000g for 1 h), membranes were collected in the 1.3M sucrose layer and at the interface with the 0.5 M sucrose layer. Purified membranes were then diluted in the HEPES buffer and centrifuged (20,000 rpm for 20 min). Thylakoid membranes were then resuspended in s sample buffer (NativePAGE, ThermoFischer) at 1mg chlorophyll. mL^-1^. Solubilization was carried out in the dark (5 min at 4°C) by adding an equal volume of 2% dodecyl maltoside. Insoluble material was removed by centrifugation (12,000 g, 20 min, 4°C) and one-tenth volume of Native PAGE 5% G 250 Sample Additive was added. Blue native PAGE was performed by using 3-12% Bis-Tris gels (NativePAGE, ThermoFischer). For two-dimensional PAGE, strips were excised and conserved at -80°C until use. After solubilization in LDS Buffer (Thermofisher) containing 5.5 M urea, two-dimensional PAGE was performed in 15% (w/v) polyacrylamide gels containing 5.5 M urea, and proteins revealed by silver staining.

### Mass spectrometry-based proteomic analysis

For proteomic analysis, 2D gels were stained by Coomassie brilliant blue and proteins were digested in-gel using trypsin (modified, sequencing purity, Promega), as previously described (Casabona et al., 2013). The resulting peptides were analyzed by online nanoliquid chromatography coupled to MS/MS (Ultimate 3000 and LTQ-Orbitrap Velos Pro, Thermo Fisher Scientific) using a 30 min gradient. For this purpose, the peptides were sampled on a precolumn (300 μm x 5 mm PepMap C18, Thermo Scientific) and separated in a 75 μm x 250 mm C18 column (PepMap C18, 3 μm, Thermo Fisher Scientific). The MS and MS/MS data were acquired by Xcalibur (Thermo Fisher Scientific). Peptides and proteins were identified by Mascot (version 2.8.0, Matrix Science) through concomitant searches against the Uniprot database (*Chlamydomonas reinhardtii* taxonomy, January 2022 download), and a homemade classical database containing the sequences of classical contaminant proteins found in proteomic analyses (human keratins, trypsin, etc.). Trypsin/P was chosen as the enzyme and two missed cleavages were allowed. Precursor and fragment mass error tolerances were set respectively at 10 ppm and 0.6 Da. Peptide modifications allowed during the search were: Carbamidomethyl (C, fixed), Acetyl (Protein N-term, variable) and Oxidation (M, variable). The Proline software (Bouyssie et al., 2020) was used for the compilation, grouping, and filtering of the results (conservation of rank 1 peptides, peptide length ≥ 6 amino acids, false discovery rate of peptide-spectrum-match identifications < 1% (Coute et al., 2020), and minimum of one specific peptide per identified protein group). Proteins from the contaminant database were discarded from the final list of identified proteins. Proline was then used to perform MS1-based label free quantification of the identified protein groups based on razor and specific peptides. The relative abundances of the different proteins were evaluated through calculation of their intensity-based absolute quantification (iBAQ, (Schwanhausser et al., 2011)) values.

### Radiolabeling of total cellular proteins

Total cellular proteins from *C. reinhardtii* were labeled with ^35^S as described previously (Ozawa et al., 2010) with some minor modifications. Cells grown photo-autotrophically in batch cultures under LL (20 μmol photons m^-2^ s^-1^) in the presence of 2% CO_2_ enriched air until reaching a cell density of 2 x10^6^ cells mL^-1^, then switched to HL (300 μmol photons m^-2^ s^-1^) for 6 h, and resuspended (at 25 μg chlorophyll. mL^-1^) in a sulfur free HSM media to induce sulfur starvation for 1 hour. During starvation light conditions were same as they were grown previously. Total cellular proteins were labeled by adding 5 μCi mL^-1^ of [^35^S] Na_2_SO_4_ (American Radiolabeled Chemicals) for 10 min. Labeling was stopped by adding 10 mM cold Na_2_SO_4_, chloramphenicol (100 μg mL^-1^) and cycloheximide (10 μg mL^-1^). Thylakoid membranes were purified by discontinuous sucrose density gradient ultracentrifugation, and radiolabeled polypeptides were separated by (12-25%) SDS-PAGE (Nellaepalli et al., 2018). After electrophoresis, the gel was dried and exposed to an imaging plate for several days and labeling was detected by using a fluorescence image analyzer (FLA7000, Fujifilm).

### Chlorophyll fluorescence and ECS measurements

Chlorophyll fluorescence measurements were carried out at room temperature on whole cells using a Dual Pam-100 (http://www.walz.com). Samples were taken from batch or PBR cultures and placed into the PAM cuvette under constant stirring. The maximal PSII yield (Φ_PSII_) was measured as the Fv/Fm ratio at a single saturation flash illumination (15,000 µmol photons m^-2^ s^-1^, 200 ms duration) upon 15-30 min dark adaptation. For ETR (Electron Transport Rate) measurement, actinic light was increased step-wise from 3 to 800 µmol photons m^-2^ s^-1^. After 2 min stabilization at each light intensity, a saturating flash (15,000 µmol photons m^-2^ s^-1^, 200 ms duration) was applied, the PSII operating yield measured as (Fm’-Fs)/Fm’ and apparent ETR calculated by multiplying the PSII yield by the actinic light intensity. 77K chlorophyll fluorescence was measured using a SAFAS Xenius optical fiber fluorescence spectrophotometer as described previously (Tolleter et al., 2011; Dang et al., 2014). PSII/PSI ratios were determined from electrochromic shifts of carotenoids (ECS) absorbance changes measurements at 520 nm using a JTS10 spectrophotometer by measuring signals induced by a saturating single turnover flash in the absence or in the presence of PSII inhibitors hydroxylamine and DCMU at final concentrations of 1 mM and 10 μM, respectively (Tolleter et al., 2011; Dang et al., 2014).

### Accession numbers

Sequence data from this article can be found at Phytozome under the following accession number: *RH22* (also named *HEL15* or *CGLD3*) as Cre03.g166650. The RNA-seq data generated in this study were deposited to the SRA database at the National Center for Biotechnological Information under the accession number PRJNA834659.

## SUPPLEMENTAL DATA

**Supplemental Figure S1.** Southern blot analysis of wild-type and *hf_0_ Chlamydomonas* mutant strain indicates a single insertion of the paromomycin resistance cassette.

**Supplemental Figure S2.** Phylogenetic analysis of DEAD-box RNA helicases proteins.

**Supplemental Figure S3.** DEAD-box RNA helicase identified in the *Chlamydomonas* genome.

**Supplemental Figure S4.** Complementation of the *hf_0_* mutant by nuclear and chloroplast transformation.

**Supplemental Figure S5.** Nucleotide sequence of the codon optimized synthetic CrRH22 used for chloroplast transformation

**Supplemental Figure S6.** Accumulation of chloroplast transcripts as analyzed by RT-qPCR in the WT and in the mutant *hf_0_* at two light intensities.

**Supplemental Figure S7.** Maximum PSII yields and PSII/PSI ratios determined in WT and *hf_0_* cells grown under LL or HL.

**Supplemental Figure S8.** Blue native PAGE of thylakoid complexes from WT and hf0 grown under HL in the absence or presence of chloramphenicol.

**Supplemental Figure S9.** Light-induced expression of CreRH22, HSP70B and HSP90C genes.

**Supplemental Figure S10.** List of PCR primers.

**Supplemental Data Set S1.** MS-based proteomic characterization of thylakoid proteins from *hf_0_* cells exposed to HL and present in the two-dimensional blue native-PAGE region of interest.

**Supplemental Data Set S2.** RNA-seq results from *hf*_0_ and wild-type control cells cultivated in low light (30 µmol photons m^-2^ s^-1^) and then exposed to high light (240 µmol photons m^-2^ s^-1^) for 2h and 24h.

## ACKNOWLEDGEMENTS

This work was supported by the French ANR (Agence Nationale pour la Recherche) projects ALGOH2, OTOLHYD and by the A*MIDEX project (n° ANR-11-IDEX-0001-02) funded by the « Investissements d’Avenir » French Government program. Experimental support was provided by the HélioBiotec platform (funded by the European Regional Development Fund, the Région Provence Alpes Côte d’Azur, the French Ministry of Research, and the “Commissariat à l’Energie Atomique et aux Energies Alternatives”). Dr. Mohamed Barakat (CNRS, CEA Cadarache) is acknowledged for managing the deposit of RNAseq files. The proteomic experiments were partially supported by Agence Nationale de la Recherche under projects ProFI (Proteomics French Infrastructure, ANR-10-INBS-08) and GRAL, a program from the Chemistry Biology Health (CBH) Graduate School of University Grenoble Alpes (ANR-17-EURE-0003).

## AUTHOR CONTRIBUTIONS

E.B.D.T. and G.P. designed the study. E.B.D.T., S.N., P.A., A.B., F.C., S.C., V.E., M.H., B.G., M.S.R., I.T., performed the experiments. E.B. performed phylogenetic and transcriptomic analysis, S.B. performed proteomic analysis. E.B.D.T., S.N., S.C., S.D., Y.C., Y.T. and G.P. analysed the data. E.B.D.T., S.N., Y.L.B and G.P. wrote the manuscript.

