## Supplementary information for "A plastidial DEAD box RNA helicase plays a critical role in high light acclimation by modulating ribosome biogenesis in *Chlamydomonas reinhardtii*"

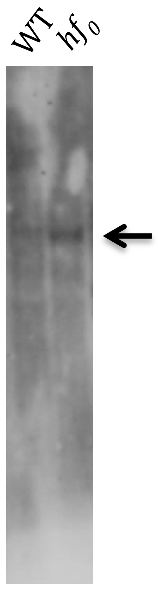


**Supplemental Figure 1. Southern blot analysis of wild-type and *hf0* *Chlamydomonas* mutant strain indicates a single insertion of the paromomycin resistance cassette.** *NcoI*-restricted genomic DNA was loaded on an agarose gel, Southern blotted and hybridized with a probe against the *AphVIII* gene (paromomycin resistance cassette).

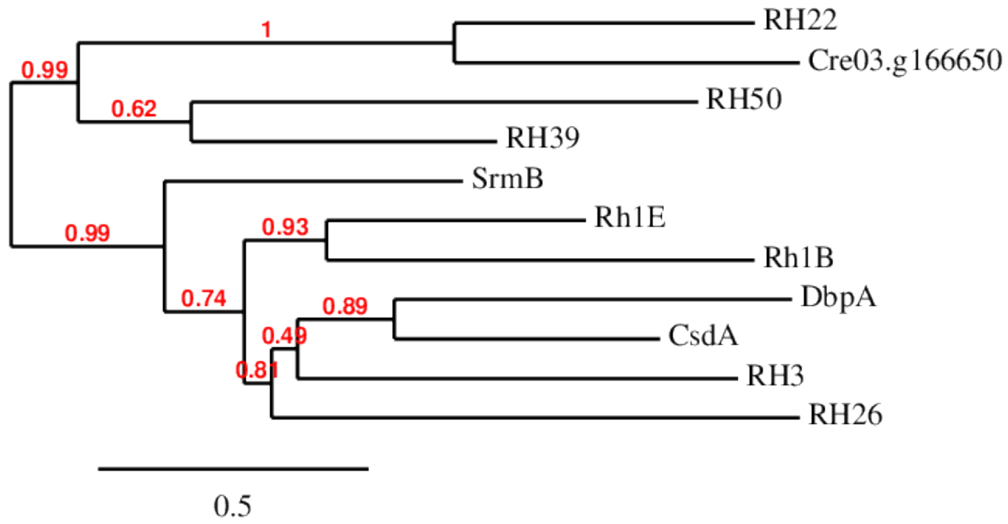


**Supplemental Figure 2. Phylogenetic analysis of DEAD-box RNA helicases proteins.** 11 homologous amino acids sequences of DEAD-box proteins from *E. coli*, chloroplast of *A. thaliana* and *C. reinhardtii* were compared by performing a phylogenetic analysis. Method used and sequences from *E. coli* and *A. thaliana* were the same as those described in (Chi et al., 2012). Corresponding GenBank accession numbers of the proteins were as follows: RH22 (NP_176207.1); RH50 (AEE74483.1); RH39 (AEE82787.1); RH26 (AED91329.1); RH3 (AED93565.1); DbpA (WP_001295553.1 replace YP_001730342.1); CsdA(DeaD) (WP_001295553.1 replace YP_001732019.1); RhlB (ADR29162.1); RhlE (WP_000007144.1 replace YP_002328316.1) and SrmB (WP_000219193.1 replace YP_002330351.1) and Cre03.g166650 for the *Chlamydomonas reinhardtii* sequence in Phytozome. The phylogenetic unrooted consensus tree was built on line with the web service Phylogeny.fr (Dereeper et al., 2008). Sequences were aligned with the Muscle program. Poorly aligned positions and divergent regions were eliminated using the Gblocks program. The tree was obtained using a Maximum Likelihood approach with the PhyML program and TreeDyn was used to visualize the tree. Bootstrap value calculations on 100 replications are indicated in red.

| Name in tree & alignment | Phytozome ID | Green cut | Gene cluster | Ribosome interactome | Predalgo |
|---|---|---|---|---|---|
| Cre-RH01 | Cre07.g319750.t1.2 | | c1 | | C |
| Cre-RH02 | g4363.t1 | | - | | O |
| Cre-RH03 | Cre17.g727700.t1.3 | | c1 | $L5^{0.01}/S3^{0.01}$ | C |
| Cre-RH04 | Cre06.g298650.t1.2 | | - | | SP |
| Cre-RH05 | Cre12.g539100.t1.3 | | c2 | | O |
| Cre-RH07 | Cre02.g118300.t1.1 | | c2 | | O |
| Cre-RH08 | Cre04.g223850.t1.2 | | 4 | | O |
| Cre-RH10 | Cre12.g505200.t1.2 | | c2 | | O |
| Cre-RH11 | Cre10.g427700.t1.2 | | c2 | | O |
| Cre-RH13 | Cre01.g022350.t1.2 | | c1 | | C |
| Cre-RH14 | Cre01.g021600.t1.2 | Hel1 | c2 | | O |
| Cre-RH15 | Cre17.g719000.t1.2 | | c8 | | O |
| Cre-RH16 | Cre12.g526850.t1.2 | | c1 | | O |
| Cre-RH17 | Cre03.g188550.t1.2 | | c14 | | O |
| Cre-RH18 | Cre07.g314900.t1.3 | | c1 | | O |
| Cre-RH20 | Cre01.g028200.t1.3 | | c14 | | O |
| Cre-RH21 | Cre13.g592150.t1.1 | | - | | C |
| Cre-RH22 | Cre03.g166650.t1.2 | Hel15 | c1 | $L5^{0.05}/S3^{0.01}$ | C |
| Cre-RH24 | Cre12.g522850.t1.2 | | - | | O |
| Cre-RH25 | Cre16.g662000.t1.2 | | c13 | | O |
| Cre-RH28 | Cre10.g420900.t1.2 | | c1 | | O |
| Cre-RH29 | Cre12.g513701.t1.2 | | c1 | | O |
| Cre-RH32 | Cre03.g156150.t1.3 | | c1 | | O |
| Cre-RH35 | g7783.t1 | | - | | O |
| Cre-RH36 | Cre16.g683500.t1.2 | | c2 | | O |
| Cre-RH38 | Cre06.g306850.t1.2 | | - | | O |
| Cre-RH39 | Cre01.g033832.t1.1 | Hel7 | - | | C |
| Cre-RH42 | Cre16.g676400.t1.2 | | - | | O |
| Cre-RH47 | Cre07.g351600.t1.2 | Hel40 | c3 | | M |
| Cre-RH47b | Cre07.g349300.t1.2 | | c1 | $L5^{0.01}/S3^{0.01}$ | M/C |
| Cre-RH50 | Cre10.g436650.t1.3 | Hel45 | c1 | $L5^{0.01}/S3^{0.01}$ | C |
| Cre-RH57 | Cre05.g238900.t1.3 | | c15 | | O |

**Supplemental Figure 3. DEAD-box RNA helicase identified in the *Chlamydomonas* genome.** Listed are DEAD-box RNA helicase genes previously identified based on homologies with *Arabidopsis thaliana* (Asakura et al., 2012). DEAD-box RNA helicases identified in GreenCut2 (Karpowicz et al., 2011) are shown in column 3. Cluster memberships as defined in (Zones et al., 2015) are shown in column 4 (among 18 clusters, c1 and c2 clusters contained genes whose expression is transiently induced in a sharp spike at the first light time point). Proteins identified as belonging to the chloroplast ribosome interactome (Westrich et al., 2021) are shown in column 5 (shown $^{0.01,\ 0.05}$ are significance of interaction between helicases and L5 or S3 ribosome subunits). Subcellular localization of *Chlamydomonas* DEAD-box RNA helicases predicted using the Predalgo tool (Tardif et al., 2012) are shown in column 6. M, C and SP correspond to targeting predictions to mitochondria, chloroplast and secretion pathway, respectively. O stands for absence of predicted targeting.

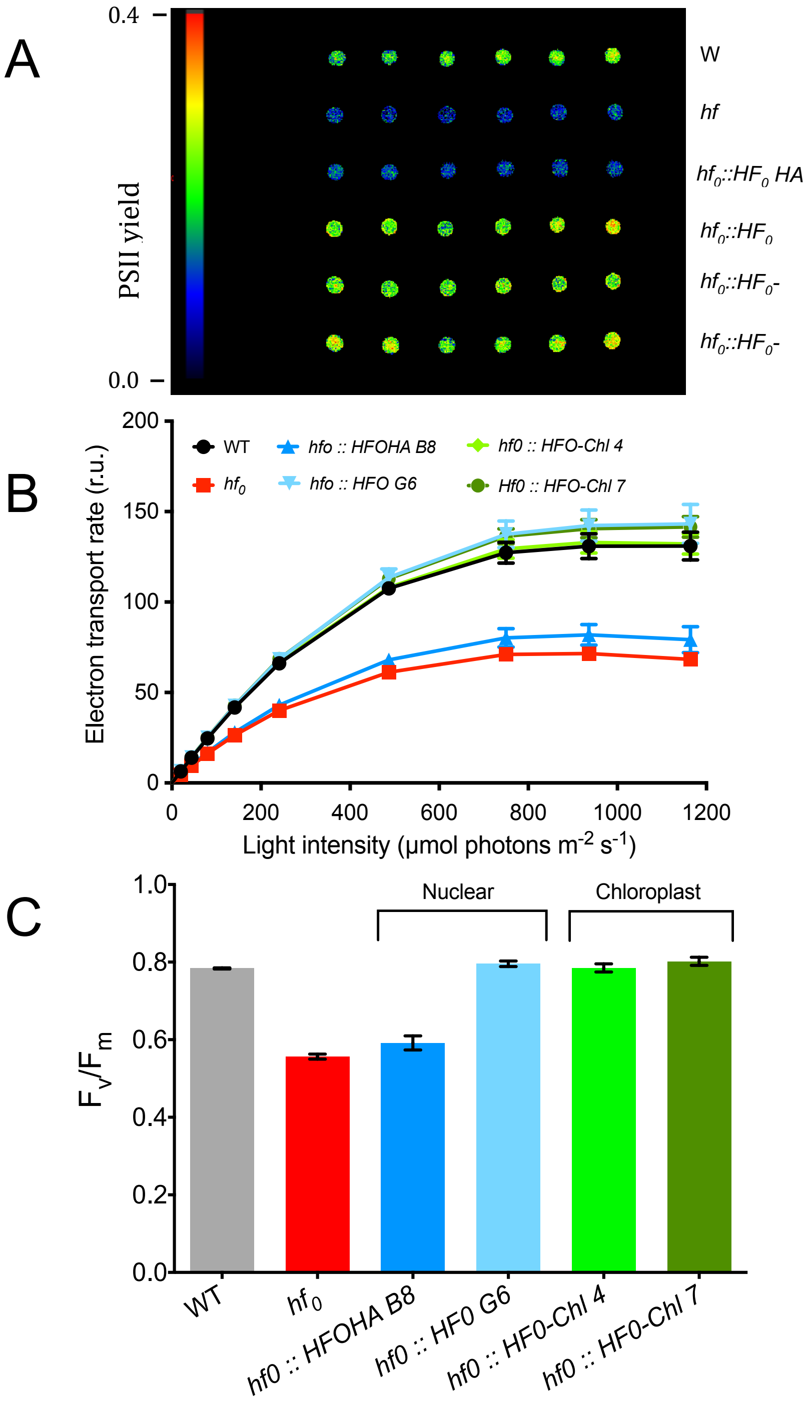


**Supplemental Figure 4. Complementation of the *hf₀* mutant by nuclear and chloroplast transformation.** (A) Antibiotic resistant clones isolated upon transformation nuclear transformation or chloroplast transformation were screened by chlorophyll fluorescence imaging. (B, C) Two nuclear transformants, two homoplasmic chloroplast transformants, and the wild-type were grown photo-autotrophically under 150 µmol photons $m^{-2}$ $s^{-1}$ before performing PAM chlorophyll fluorescence measurements. (B) Electron transport rates estimated as PSII yields x incident light intensity). (C) Maximal PSII yield (measured as $F_v/F_m$). Full complementation was observed in one nuclear transformant and in two independent chloroplast transformants.

```
ATGCGTGCTTTACCACGTACAACAGGTCCACAAGCTGGTCGTCAAACATCACCAACACATGTTCC
AATTGCTACACATGCTGTTTCACCATTAAATTGTGCTCATGCTTGGTGGCGTCCAGTTGGTGATG
TATCACGTTCATCAGTTGCTGGTTCATTTGCTGCTGGTTCAACACGTGCTTCACGTCCACCACGT
GTTGCTGTTTCAGCTTGTTTATCAACATGGGCTCCAACACCATCTGATGCAGGTTTATTATCAGG
TGAAGAAGCTTTCTTTGCTTCAGAACATGCTTCATTCGCTGAATTAGGTGTTTCACCAGCTATTC
AAGCTGCTTTAGAAACATCAGGTTTAAAACGTCCATCACGTATTCAAGAATTAGCTGCTCCACAT
ATTTTACGTGGTCGTAATGTTGTTGTTGCTGCTGAAACAGGTTCAGGTAAAACATTATCATATTT
AGTACCAATTGCTCACTTAATGTTAAAACAACGTACAGCTATGCAACAACACGCTGCTGCATTAG
AAGCTGCTGGTCCAGCTGCTGATGGTGCTGAACGTCCACGTTTTACACGTTATTTAGCTTTAGTT
TTATGTCCAAATGTTGCTTTATGTCAACAAGTTGCTGCTGCTGTTAATGCTTTACGTGGTCCTGC
TGCTGCAGATGGTTCATCTCAACAACAACAACAACAACAATTAGTTTCAGCTGCTGTTATTAACT
CATCAAACCCACCACCATTTGAAACACCAGATGTTGTTGTAGCTACACCAGCTGGTTTATTAAAT
ATTATTGATGACGCAGGTGGTGCTTACGGTTGGTTATGGTCTGAAGAAGGTATGCAAGCTCGTAT
TCGTCACGTTGTTTTAGACGAAGCTGATTTATTATTAACACCAGCTTATTCACGTGCTACACAAC
GTATTTTAACATTATTCAAAACAGCTGATCGTCGTCGTGTTGAAGCTAAATTATTTGATGAATTA
GGTTTAGCTGGTAAAGATGAATTTGATCGTTTACCACGTGCTTTACAAGTAGCTGCTTGGACAGG
TGGTGCTCCAGCTATGTTAGCTGAAGGTTATCGTCCAAAACGTTCATTAAACCCAGATGCTAAAT
ATGGTCCATATTGGCGTCGTCAATACATTTATTCAGCTGCTACATTACCAGCTGCTACTTATTCA
GATGTTGGTTCAGCTATTGCTAAAGCTCATCCAGATGCTGTTTGGGTTTCAACAGACTTATTACA
TTCATCAAAACCACAAGTTGAACACGCTTGGCGTGAAGTTCGTGATGATGATTTTACAACAAACT
TATTAGATTCTATTAAATCAGATCCAGATTATCAAGCTCGTTCTGGTAAAACTTTAGTTTTTGCT
GCTGACGGTGCTTCAGCTGATGCTGCTTCTGAAGTTTTAGCAGGTGGTAATGTTCCACACGTTGT
TTATCACAAATCACGTCCAATGGGTGAACAAGCTGCTGCTTTAGCTACATTACGTGAACAACCAG
GTTGTGTTATGGTTTGTACAGATGCTGCTGCTCGTGGTTTAGATGTTGAAGATATTAAACACGTT
GTTCAAGCTGATTTCGCTGCTAACGCTATTGATTTCATTCATCGTATTGGTCGTACAGCTCGTGC
TGGTCGTGGTGGTCGTGTTACATCATTATATCGTGAACACAACCGTGCTTTAGTTGAAGTTTTAC
AATCATATATTGCTGATGGTGTTGCTTTAGAAGCTGCATTTTCACGTGCTCGTTCATTTTCAAAA
AAATTAAAAAAAACAGGTGGTGTTTTTGTTCCACGTGGTATGGCTGCAGGTGGTGAAGAAGCTGA
AGCTGCTCAAGCTAATGCTGGTGAAGGTGCAGGTGAAGGTCAAGAAGGTGCTGCTGCAGGTCCAG
CAGCAGGTCGTGGTGGTCCAGGTCGTGGTGCAGGTGGTGGTCGTGGTGCAGGTCGTGGTCGTGGT
CGTGCTGGTCGTGAACAAGCTGCTCAAGAAGAAGCTGTTCGTTCATAA
```

**Supplemental Figure 5. Nucleotide sequence of the codon optimized synthetic CrRH22 used for chloroplast transformation.**

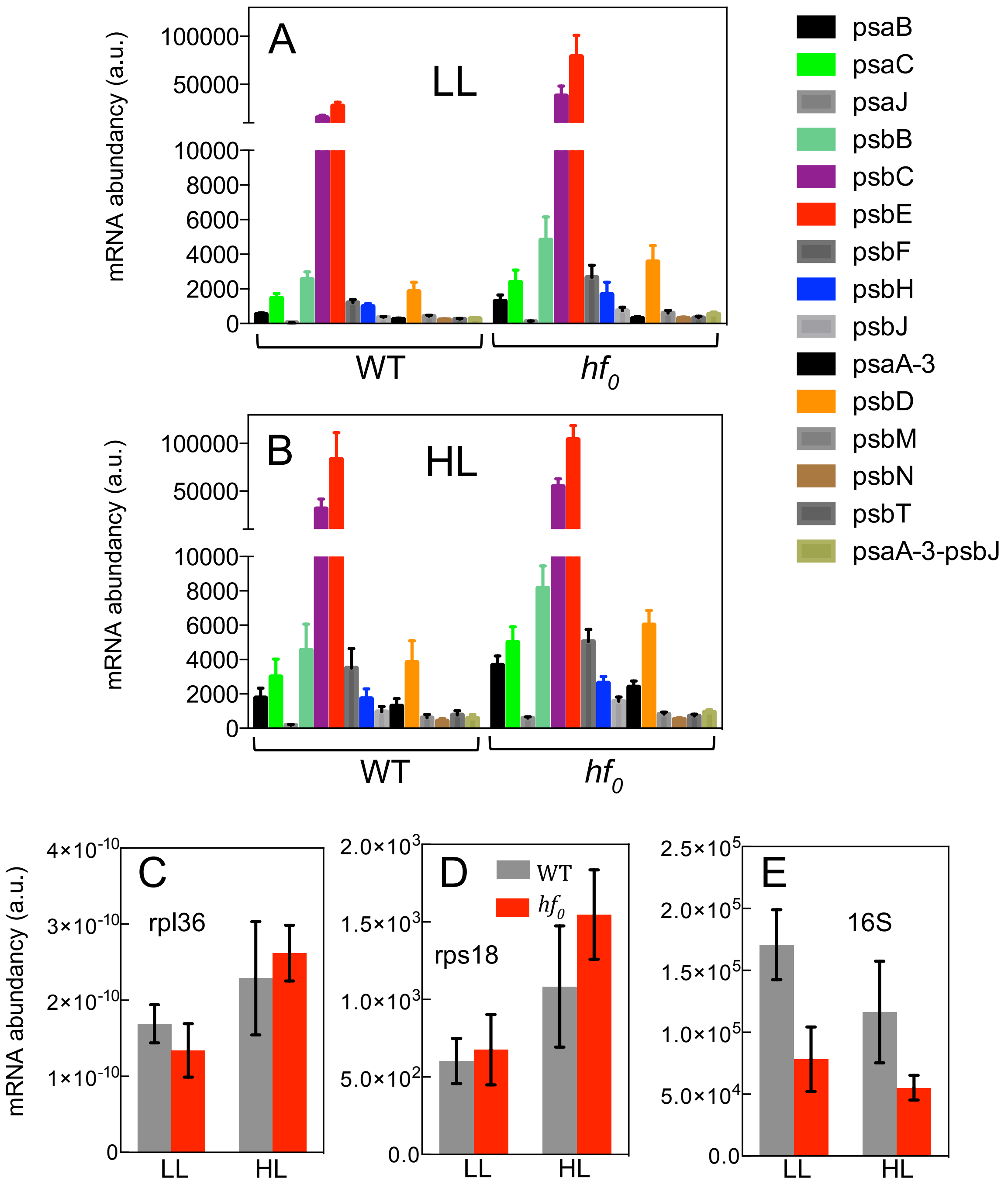


**Supplemental Figure 6. Accumulation of chloroplast transcripts as analyzed by RT-qPCR in the WT and in the mutant *$hf_0$* at two light intensities.** (A, B) Relative transcript levels of different plastidial genes encoding subunits of PSI and PSII under (A) LL (30 µmol photons $m^{-2}$ $s^{-1}$) or (B) HL (240 µmol photons $m^{-2}$ $s^{-1}$). Data were normalized on *rpl36* ribosomal subunit expression levels. (C, D) *rpl36* and *rps18* transcript levels are not affected in the *$hf_0$* mutant as compared to the WT. (E) 16S rRNA transcripts show a 2-fold reduction in *$hf_0$* as compared to the WT in both light conditions. Shown are means of three technical replicates ± SD.

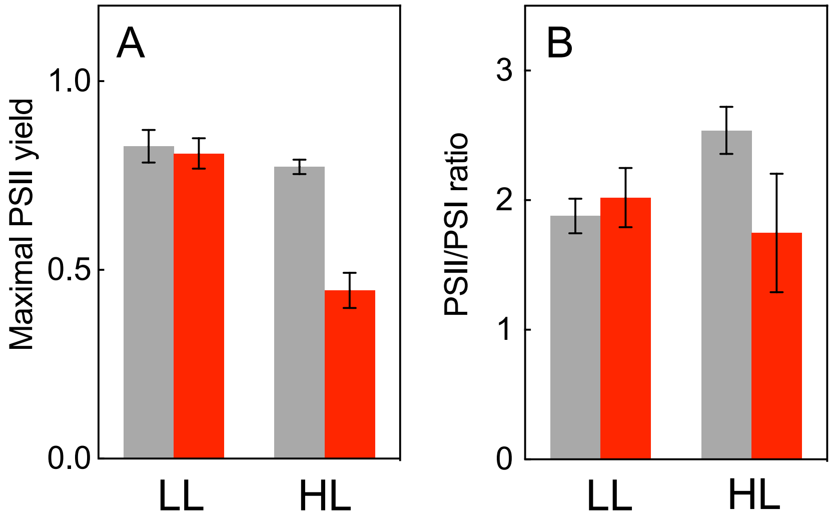


**Supplemental Figure 7. Maximum PSII yields and PSII/PSI ratios determined in WT and *hf₀* cells grown under LL or HL.** (A) Maximal PSII yield was determined chlorophyll fluorescence measurements as $F_v/F_m$. (B) PSII/PSI ratios were determined from ECS absorption measurements in the absence or in the presence of PSII inhibitors 10mM hydroxylamine and 10μM DCMU. Shown are means of biological replicates ± SD (n=3).

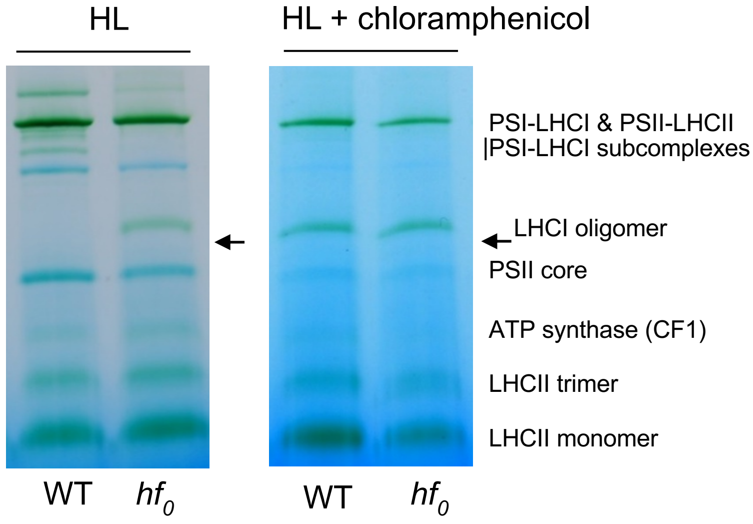


**Supplemental Figure 8. Blue native PAGE of thylakoid complexes from WT and *hf₀* grown under HL in the absence or presence of chloramphenicol.** WT and in *hf₀* cells were grown photo-autotrophically in photobioreactors under LL and then switched to HL. The two left panels (same as on Fig. 7F) are in the absence of chloramphenicol. The two right panels are in the presence of chloramphenicol (50 mg. $L^{-1}$), added to the culture 1h before the HL switch. Arrow shows the green band identified as appearing in *hf₀.* after HL exposure. The same band appears in both strains in response to the chloramphenicol treatment.

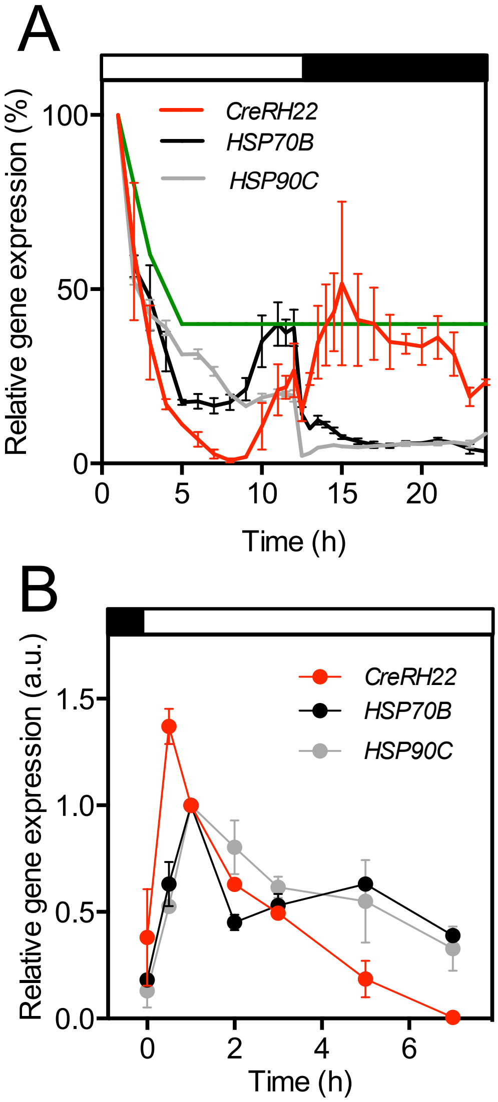


**Supplemental Figure 9. Light-induced expression of *CreRH22*, *HSP70B* and *HSP90C* genes.** (A) Gene expression data have been obtained from (Zones et al., 2015). In this study, *C. reinhardtii* cells were grown in day-night cycles. The continuous green line shows the light stress cluster threshold as defined in (Zones et al., 2015). (B) The relative expression of *CreRH22*, *Hsp70B* and *HSP90C* was followed by qPCR during a day-light (200 µmol photons $m^{-2}$ $s^{-1}$) transient. Data were normalized on *RACK1* transcript levels. Shown are mean ± SD of two independent experiments.

| Primer name | Sequence (5' to 3') | Use |
|---|---|---|
| GW Aph8 Fw1 | CTGGTGCTGCGCGAGCTGGCCCACGAGGAG | Genome walker |
| GW Aph8 Fw2 | TGGTTCGGGCCGGAGTGTTCCGCGGCGTT | Genome walker |
| GW Aph8 Rev1 | CCAGCGCGAGATCGGAGTGCCGGTCCG | Genome walker |
| GW Aph8 Rev2 | CGAGACTGCGATCGAACGGACACCGC | Genome walker |
| AP1 | GTAATACGACTCACTATAGGGC | Genome walker |
| AP2 | ACTATAGGGCACGCGTGGT | Genome walker |
| aph ORF For | CGAAGCATGGACGATGCGTT | Southern |
| aph tail3 | CGAGACTGCGATCGAACGGACA | Southern |
| CDS helicase Fw1 | ATGCGGGCGCTGCCACGGACGA | Nuclear expression |
| CDS helicase Rev1 | TCACGACCGCACCGCCTCCTC | Nuclear expression |
| helicase cpt BglII Fw | AGATCTCACCATGTGCGTGCTT | Chloroplastic expression |
| helicase cpt BglII Rev | AGATCTTGAACGAACAGCTTCT | Chloroplastic expression |
| chel F2 | CCATGCTCGCGGAGGGCTACC | CrRH22 expression |
| chel R | CTGCGTCCGTGCACACCATCAC | CrRH22 expression |
| 5'Rack | CTTCTCGCCCATGACCAC | Nuclear housekeeping gene |
| 3'Rack | CCCACCAGGTTGTTCTTCAG | Nuclear housekeeping gene |
| qRPL36F | AAGGGCCATGTGGTCACTAA | Chloroplastic housekeeping gene |
| qRPL36R | TACAGCGACTTACCCCCTTG | Chloroplastic housekeeping gene |
| qpsaA-3F | TGGGGTACGGTTACAGCTTC | RTqPCR |
| qpsaA-3R | CGAGAATGCCCATACGAAGT | RTqPCR |
| qpsaBF | TATTGGCCTTGGTGACTTCC | RTqPCR |
| qpsaBR | CGTCACATGGGAAGCTGTAA | RTqPCR |
| qpsaCF | CCATGGGATGGTTGTAAAGC | RTqPCR |
| qpsaCR | CACAGTCTTCAGTGCGTGGA | RTqPCR |
| qpsaJF | CGTGGTGGTCGAGTTAAAGA | RTqPCR |
| qpsaRF | CAGCTGTTTTTGACCCCATT | RTqPCR |
| qpsbBF | CCCACGTGGTTGGTTTACTT | RTqPCR |
| qpsbBR | TGCCAAATGTGACCAAAGAA | RTqPCR |
| qpsbDF | GTTTATGGGCATTCGTTGCT | RTqPCR |
| qpsbDR | CTGACGAAGCATGAAACCAA | RTqPCR |
| qpsbFF | TCCTATTTTCACAGTTCGTTGG | RTqPCR |
| qpsbFR | CGTTGAATGAATTGCATAGCAG | RTqPCR |
| qpsbHF | CACTGGCCTTCCGTTAAGAT | RTqPCR |
| qpsBHR | TCAGGCAATTTGCTTACACC | RTqPCR |
| qpsbJF | TCCCTCTATGGCTTGTTGGT | RTqPCR |
| qpsbJR | TGCGCCAATAGCTAAAGTACC | RTqPCR |
| qpsbMF | TTGTTCACGCTTATGTCTTCGT | RTqPCR |
| qpsbMR | TCCCTTTCTGTATAACCGCTAA | RTqPCR |
| qpsbNF | GGCACCGAAGTCACTTGTAAA | RTqPCR |
| qpsbNR | TGATCCTTTTGAAGAACACGAA | RTqPCR |
| qpsbTF | CGATTCTTAACAGCCCCAGA | RTqPCR |
| qpsbTR | TTAATAGACCTTTTGCCGTTG | RTqPCR |
| q intergene psaA-2/psbJ F | AGTCTTCGGAGGAATGCAAA | RTqPCR |
| q intergene psaA-2/psbJ R | GCTATTCCCTTTCAGGTCCA | RTqPCR |
| q16SF | TGGAGGAAGGTGAGGATGAC | RTqPCR |
| q16SR | TCACGAGTCGCTTCTGATTG | RTqPCR |
| qrps18F | CACGTTTAACAGCAAAACAACAA | RTqPCR |
| qrps18R | CAAACGGTAACAAGCCCATT | RTqPCR |
| 16S 23F | TCCATGGAGAGTTTGATCCTG | Northern |
| 16S 300R | TCCTCTCAGACCAGCTACTGC | Northern |
| 7S sde F | GAATTAAGGCGTACGGTGG | Northern |
| 7S sde R | CCCACGACCCCTACATC | Northern |
| 3S sde F | CGAAGCAGCTGAATCCTGCAC | Northern |
| 3S sde R | CTACTGGACTTTCACCATCTATGG | Northern |
| psbD sde F | GTTTATGGGCATTCGTTGCT | Northern |
| psbD sde R | TGAAACGGAAGATAGCAGC | Northern |
| psaA-2 sde F | TAAACCAGGACATTTTTCACG | Northern |
| psaA-2 sde R | CCCACTTAACCAAATGAAAATG | Northern |

**Supplemental Figure 10. List of PCR primers.**
